# A Serpin27A-dependent Toll Signaling Underlies Host Genetics of Cancer Resistance in *Drosophila*

**DOI:** 10.1101/2021.03.13.435228

**Authors:** Reeta Singh, Sneh Harsh, Anjali Bajpai, Subhabrata Pal, Ravi Kant Pandey, Pradip Sinha

## Abstract

Cancer resistance varies amongst individuals, although its host genetic underpinnings remain largely elusive. Remissions of sarcomas were first reported following repeated injections of patients with mixtures of killed bacteria—Coley’s toxins—a phenomenon, which was subsequently causally traced to induction of innate immunity. Here we reveal remission of *Drosophila* epithelial neoplasms by genetically triggered host innate immunity via Toll signaling. These neoplasms display capacities to receive and, in rare instances, induce Toll signaling. A tumor-induced and progressive Toll signaling, however, did not culminate in tumor suppression. By contrast, *Drosophila* hosts heterozygous for *spn27A*^*1*^ mutation, which constitutively produce activated Toll ligand, Spz^Act^, displayed comprehensive tumor remission via Toll-induced, NF-κB-mediated, tumor cell death. Our results reveal a novel node of host genetic cancer resistance via serpin-dependent Toll signaling.

## Main text

Towards the end of the 19th century, when William Coley observed remissions of sarcomas in patients by repeated injections of killed cocktails of bacterial strains—subsequently termed as Coley’s toxins (Nauts et al., 1953)—it gave birth to the field of cancer immunotherapy: more specifically, innate immunity-mediated cancer remission. Toll signaling, a core and a highly conserved innate immune pathway (Rakoff-Nahoum and Medzhitov, 2009), thus holds the promise of unraveling some of the elusive (Klein, 2009) host genetic underpinnings of cancer resistance. Genetic tractability and conservation of Toll signaling in *Drosophila* makes it an ideal model organism to probe these possibilities.

In *Drosophila,* Toll signaling is triggered by its activated ligand, Spz^Act^, via cleavage of its precursor zymogen, pro-Spz, by serine proteases (SPs). SPs remain inactive when bound with their negative regulators, Serpins (De Gregorio et al., 2002; Hashimoto et al., 2003). Downregulation of serpins activate SPs and cleavage of pro-Spz zymogen to activated ligand, Spz^Act^; its binding with the Toll receptor then activates its nuclear targets in a tissue context dependent manner via one of the *Drosophila* homologs of mammalian NF-κB transcription factor, namely, Dorsal (Dl), Dorsal immune-like factor (Dif) or Relish (Rel) (Lemaitre et al., 1996; Meyer et al., 2014). These nuclear targets of Toll signaling include ventralizing gene sets in embryos (Hashimoto et al., 2003), proapoptotic genes *grim, reaper or hid* in the epithelial cells (Meyer et al., 2014), or anti-microbial peptide coding genes, such as *drosomycin (drs)* in larval or adult fat body (Lemaitre et al., 1996). Expression of a *drs-GFP* transgene in larval or adult fat body serves as a sensor of systemically produced Spz^Act^ from the site of injury or microbial infection (Lemaitre and Hoffmann, 2007). During microbial infection, its expression is initially seen in the posterior adipocytes of the larval fat body (*drs-GFP,* **Figure 1A**), which subsequently propagates to the anterior adipocytes (*drs-GFP*, **Figure 1B)** aided by Spz^Act^ derived from the circulating host hemocytes (Shia et al., 2009) (**Figure S1**). Thus, knockdown of *spz* in host hemocytes abrogates propagation of *drs-GFP* expression without affecting its initiation in fat body by the Spz^Act^ ligand (**Figure 1C D)**.

**Figure 1.**
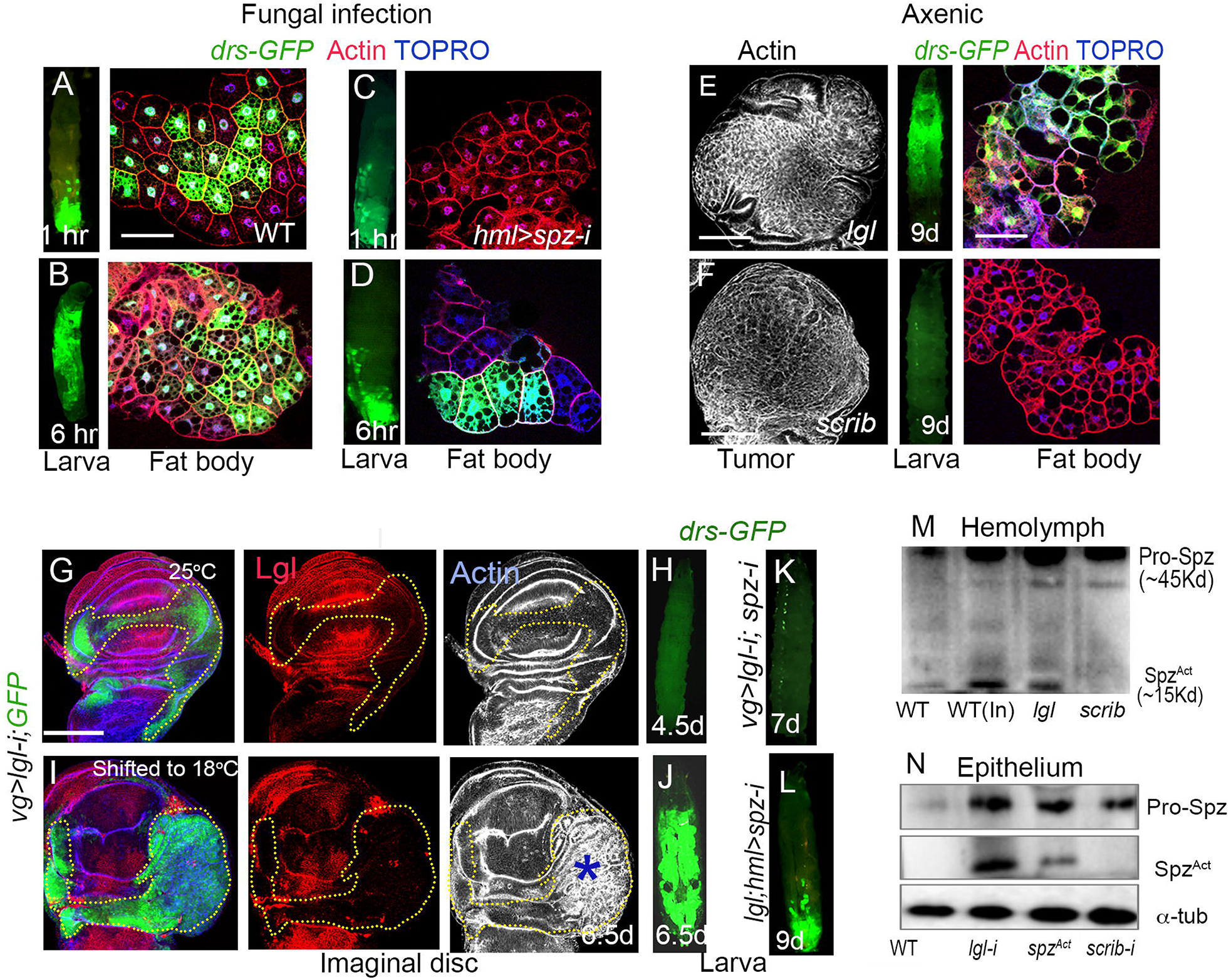
Spz^Act^ from *lgl* neoplasms initiate host Toll signaling, which is propagated by its hemocytes. (**A-D**) Initiation (1 hr, A) and propagation (6 hr, B) of systemic Toll signaling (*drs-GFP*) in fat body of wild type larvae following fungal infection. Knockdown of *spz* in hemocytes (*hml>spz-i*) in these larvae did not affect initiation (*drs-GFP*, C) but blocked propagation of Toll signaling (*drs-GFP*, D). (**E, F**) *lgl* larva (E), but not that of *scrib* (F) displays Toll signaling without microbial infection (axenic). Explanted fat body larvae in (A-F) are shown at higher magnification to reveal their *drs-GFP* (green) expression. (**G-J)** Wing imaginal disc displaying loss of Lgl (red, dotted line) upon knocked-down *lgl (vg>lgl-i*) in wing imaginal discs of larva grown at 25°C (G); *drs-GFP* (green) in host larval fat body is not induced (H). Upon shift of larval cultures shown in (G) to 18°C after that prolongs larval life, these imaginal discs displayed neoplastic transformation (star, actin, I) and robust *drs-GFP* (green) expression in host larval fat body (J). (**K, L**) Wild type larva upon simultaneous knockdown of *spz* and *lgl* in their wing imaginal disc epithelium fail to trigger *drs-GFP* (green) reporter expression (K). *lgl* larva displaying knockdown of *spz* from their hemocytes displayed drs-GFP in posterior adipocytes their fat body (*drs-GFP*) but failed to propagate its expression anteriorly. (**M, N**) Western blots of larval hemolymph (M) or epithelial neoplasms (N) to detect the activated Toll ligand, Spz^Act^. All larval images are displayed at the same magnification. Larval ages in this and subsequent figures are shown in <u>d</u>ays <u>a</u>fter <u>e</u>gg <u>l</u>aying (dAEL). Scale bar=100μm

Cancer patients often display sterile inflammatory responses, including Toll signaling (Rakoff-Nahoum and Medzhitov, 2009). To test if *Drosophila* tumors, too, trigger Toll signaling, we examined expression of the *drs-GFP* reporter in *lgl* and *scrib* mutant larval fat body, while their imaginal disc epithelia displayed neoplastic transformations due to loss of Lgl (Agrawal et al., 1995) and Scrib (Bilder and Perrimon, 2000) tumor suppressors, respectively. We noticed *drs-GFP* expression in fat body of *lgl* (**Figure 1E**) but not in that of *scrib* (**Figure 1F, Figure S2**) larvae; the latter observation is contrary to a previous claim (Parisi et al., 2014). We also noticed that *Drosophila* larval immune organs like the fat body and lymph glands express Lgl (**Figure S3**). Toll signaling seen in the *lgl* larvae could thus be fallouts of loss of Lgl from these immune organs, rather than a systemic trigger from their epithelial neoplasms. To allay this concern, we induced selective epithelial loss of Lgl using a *vg-Gal4* drive; the latter expresses only along the D/V margins of the developing wing and haltere imaginal discs, besides larval salivary glands—leaving out its immune organs (**Figure S3**). At 25°C larval growth temperature, loss of Lgl in wing epithelium (*vg>lgl-RNAi*) neither displayed neoplastic transformation (actin, **Figure 1G)** nor induced Toll signaling (*drs-GFP,* **Figure 1H)**. However, upon shift of these larval cultures to a lower growth temperature of 18°C—following 4.5 days of development at 25°C—and a consequent prolongation of larval life by two additional days (6.5 d AEL)—we noticed a robust neoplastic transformation in the wing epithelium (**Figure 1I**) besides induction of *drs-GFP* expression in the larval fat body (**Figure 1J**). These results therefore revealed induction systemic Toll signaling by *lgl* neoplasms. Of note, these *lgl* neoplasms also express Spz (**Figure S4**) thereby suggesting their production of tumor-derived Spz^Act^— reminiscent of the fallout of its gain in a wild type imaginal epithelium (**Figure S4**). We also noted that clonal progression *lgl* neoplasms was marked progressive increase in the numbers of *drs-GFP* expressing adipocytes, commensurate with the buildup tumor-derived Spz^Act^ (**Figure S5**). Not surprisingly, knockdown of *spz* in *lgl* neoplasms extinguished their systemic Toll signaling altogether (**Figure 1L**) while its knockdown in host larval hemocytes (**Figure 1K**) or genetic ablation of the latter (**Figure S5**) selectively abrogated propagation of the tumor-triggered Toll signaling to the anterior adipocytes.

By contrast to *lgl* neoplasms, those induced by individual knockdowns of Scrib (Bilder and Perrimon, 2000), Avalanche, Avl (Colombani, 2012) or Rab5 (Morrison et al., 2008) tumor suppressors did not induce systemic Toll signaling, although their neoplasms displayed expression of the *spz-GFP* reporter (**Figure S6**). Western blots of hemolymph proteins from fungus-infected wild type or uninfected *lgl* larvae displayed Spz^Act^, unlike that of uninfected *scrib* larvae, while all of these displayed expressions of the pro-Spz zymogen (**Figure 1M**). Further, Spz^Act^ was also seen in western blots *lgl* neoplasm proper, or in wild type epithelia expressing a *spz*^*Act*^ transgene, but not in *scrib* neoplasms (**Figure 1N**). In essence, while all the *Drosophila* epithelial neoplasms tested here produce the pro-Spz zymogen, its cleavage into an activated ligand, Spz^Act^, was seen only in *lgl* neoplasms, which reveal the causal underpinning of their induction of host Toll signaling.

To obtain clue(s) to selective induction of Toll signaling in *lgl* larvae, we examined transcriptomes of *lgl* and *scrib* neoplasms of comparable larval age. Gene Set Enrichment Analysis (GSEA) displayed significant enrichment of Toll pathway in *lgl* neoplasms (**Figure 2A**) while transcriptome of *scrib* neoplasm, revealed a relatively modest enrichment of its Toll signaling pathway (**Figure 2B).** A closer examination of transcript levels of Toll pathway members such as genes coding for Toll receptors, SPs, Serpins and downstream effectors (**Figure S7**) revealed there near comparable expressions in these two neoplasms, which did not offer a ready explanation of selective production of Spz^Act^ in *lgl* neoplasms. We further examined *lgl* transcriptomes at different time intervals of *lgl* tumor progression. Here, we noticed a conspicuous but short-lived down-regulation of genes coding for negative regulators of SPs: for instance, a serpin, *spn27A,* early during *lgl* tumor growth (4.5 d AEL, **Figure 2C**). Further, at this stage of *lgl* tumor progression, we also noticed upregulation *spz* transcripts—that was consistent with synthesis of pro-Spz zymogen—besides a variety of SPs: for instance, *easter* (*ea*), *persephone* (*psh*), *Spz processing enzyme* (*SPE*) *or pro-phenol oxidase (PPO*) (**Figure 2C** also see **Figure S8**). These features of transcriptomes of *lgl* neoplasms, thus offered a compelling explanation of their production of Spz^Act^ (see **Figure 1**) as a net fallout of upregulation and activation SPs and down-regulation of their negative regulator, *spn27A* (**Figure 2C, S6**).

**Figure 2.**
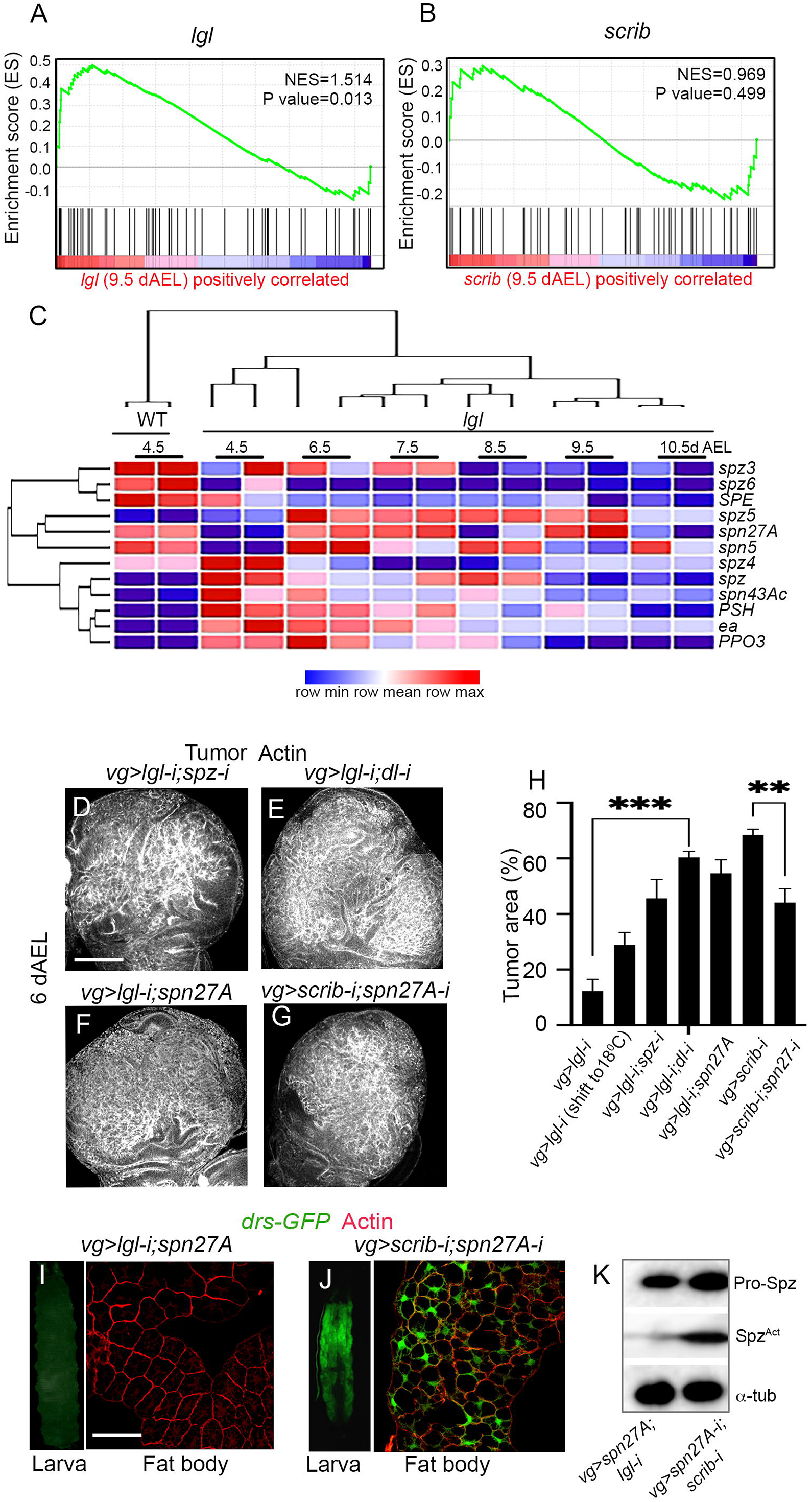
Tumor-autonomous Toll signaling retards its progression. **(A-C)** GSEA plots depicting enrichment of Toll pathway in transcriptomes of *lgl* (A) and *scrib* (B) neoplasms explanted from comparable larval age (9.5 days); y axis displays enrichment score (ES) and x axis displays ranked order of genes in descending order of their fold-changes in tumor versus wild type discs. (C) Hierarchical-clustered heat maps displaying expression statuses of genes of select Toll pathway members in *lgl* tumors with progressive larval ages (4.5-10.5d AEL). **(D-H**) Comparison of sizes of neoplasms displaying simultaneous knockdown of *lgl* and *spz* (D)*, lgl* and *dl* (E), or *lgl* knockdown with accompanying gain of *spn27A* (F) and, finally, simultaneous knockdown of *scrib* and *spz* (G); quantifications of their tumor sizes (H). (**I-K**) Gain of *spn27A* in *lgl* (I) and its knockdown in *scrib* (J) neoplasms, respectively, suppresses and induces Toll signaling (*drs-GFP*) in host larval fat body; their western blots to detect Spz^Act^ (K). Paired *t* test using GraphPad **p value= 0.0015, ***p value=0.0007. Scale bar=100μm.

What could be a tumor-autonomous impact Toll signaling in *lgl* neoplasms? We noticed that sizes of *lgl* neoplasms were initially smaller than their *scrib* counterparts (**Figure S9**). This could be due to induction of cell death by Toll signaling (Meyer et al., 2014) as seen upon gain of Spz^Act^ in imaginal disc epithelium **(Figure S9)**. Cconsistent with this rationale, we noticed that loss of Toll signaling in *lgl* neoplasm via knockdowns of Toll ligand production (*vg>lgl-i*, *spz-i;* **Figure 2D)** or loss of transduction of nuclear signal (*vg>lgl-i*, *dl-i,* **Figure 2E**) or, conversely, via overexpression of *spn27A* (**Figure 2F)** promoted their growth, overriding their staggered tumor progression (**Figure 2H**). Likewise, fain of Toll signaling via knockdown of *spn27A* in *scrib* neoplasms reduced their sizes (**Figure 2G, H**). These conditions for promotions of *lgl* tumor progression or restraint on *scrib* tumor growths were readily reconciled, respectively, with suppression of (**Figure 2I**) or induction of (**Figures 2J**) their Spn27A-dependent Spz^Act^ production (**Figure 2K**).

An initial Toll-induced and staggered progression of *lgl* neoplasms of imaginal discs, however, is overridden with increasing larval age when these attain massive sizes, which accompany their mutual invasions (**Figure S9,** also see (Gateff, 1978). We reasoned that remission of *Drosophila* epithelial neoplasms may require induction of robust Toll signaling in the host larvae—as in in patients receiving Coley’s toxin, for instance (Karbach et al., 2012; Nauts et al., 1953). Previously, it was shown that a haplo dose of a serpin coding gene, *spn27A*, due to heterozygosity of one of its null allele, *spn27A*^*1*^ brings about constitutive Toll signaling via SP-mediated cleavage of pro-Spz to Spz^Act^ (Hashimoto et al., 2003): its hemolymph being a rich source of Spz^Act^ (also see **Figure S10**). We thus examined progression of *scrib* neoplasm in a *spn27A*^*1*^/*+* host genetic background. Unlike the s*crib* neoplasms induced in a wild type genetic background (**Figure 3A**), those induced in a *spn27A*^*1*^/*+* host genetic background displayed extensive caspase activation leading to tumor elimination (**Figure 3B**), while the host larvae escaped pupal lethality—unlike their wild type counterparts (**Figure 3A**)— and emerged as adults despite wing deformities (**Figure 3C**), displaying a substantially restored adult life span (**Figure 3D**). Comparable outcomes were also seen when *lgl* neoplasms were induced in *spn27A*^*1*^/*+* host larvae (**Figure S11**). Tumor remission in *spn27A^1^/*+ hosts persisted even after knockdown of *spz* in *scrib* (**Figure 3E**) or in *lgl* neoplasms (**Figure S11**) revealing thereby that production of the Toll ligand from neoplasms *per se* is inconsequential for host genetic remission of tumors. By contrast, blocking nuclear transduction of Toll signal by knocking-down expression of Dl or Dif transcription factor in *scrib* (**Figure 3F, G**) or in *lgl* (**Figure S11**) neoplasms arrested their host genetic remission. Finally, we show that *scrib* somatic clones induced in wing imaginal discs, or *scrib* neoplasms induced by its knockdown in the developing eye and head imaginal epithelia were also eliminated in *spn27A*^*1*^/*+* host larvae (**Figure S12**). A robust and constitutive host genetic Toll signaling, therefore, is necessary and sufficient for the remission of epithelial neoplasms in *Drosophila*. We also note that host genetic cancer remission revealed here is essentially prophylactic since *serpin* mutant hosts display constitutively activated Toll signaling (Levashina et al., 1999), ahead of tumor initiation.

**Figure 3.**
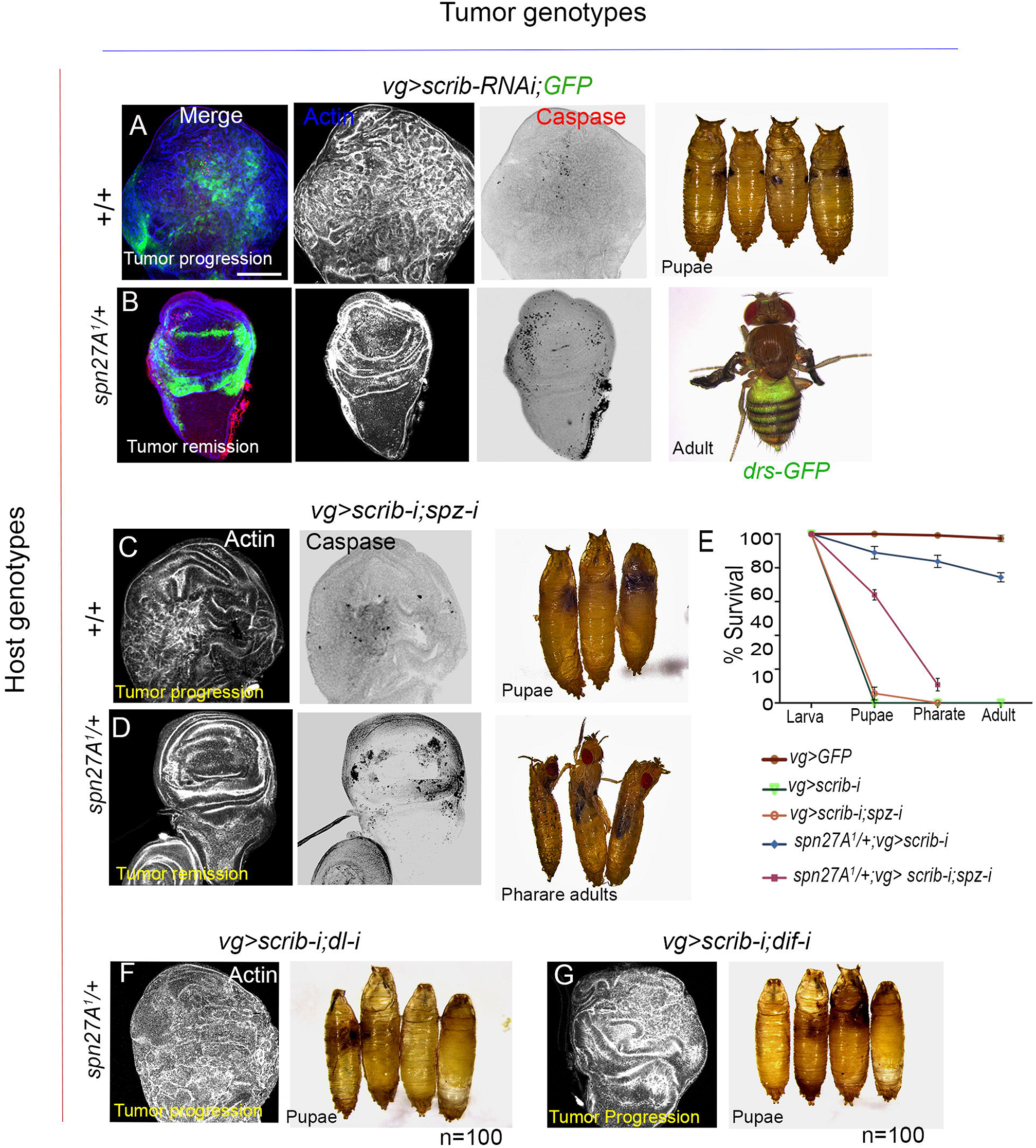
Toll-mediated remission of *scrib* neoplasms in *spn27A*^*1*^/*+* host genetic background. (**A**-**E**) *scrib* wing imaginal disc neoplasms (actin) display rare cell death (caspase) in wild type host larvae, which die as pupae (A). Conversely, *scrib* neoplasms in s*pn27A*^*1*^/*+* larval hosts display extensive cell death (caspase), tumor remission and survive as adults, d*rs-GFP* reporter expression its adult fat body (B). *scrib* neoplasms displaying knockdown of *spz* displayed rare cell death (caspase) in wild type host larvae, which died as pupae (C), while these neoplasms are eliminated by cell death (caspase) in s*pn27A*^*1*^/*+*larval hosts that survived till pharate adult stage; survival plots (E) of the animals shown in (A-D). (**F, G**) Restoration of tumor progression upon blocking reception of Toll signaling (*dl-i,* F or *dif-i,* G) in *scrib* neoplasms induced in s*pn27A*^*1*^/*+* larval hosts. Under both these conditions, host animals displayed 100% pupal lethality (n (3) =100 each set). Scale bar=100μm.

Previously, metabolically perturbed cells in *Drosophila* were shown to be eliminated by a localized Toll-mediated cell competition (Alpar et al., 2018; Meyer et al., 2014) or via systemic production of the Spz^Act^ due to microbial infection (Germani et al., 2018). Tissue surveillance mechanisms like cell competition maintain cell fitness within an organ (for recent review, see Baker, 2020) and are also effective for tumor elimination (Agrawal et al., 1995; Katsukawa et al., 2018; Khan et al., 2013). While results presented here may appear reminiscent of Toll-mediated cell competition, we do notice an important distinction: namely, that local interactions of cell neighbors—as in localized Toll-mediated tissue surveillance (Alpar et al., 2018; Meyer et al., 2014)—may not suffice for tumor elimination, particularly when the latter displays sufficient growth. Instead, the paradigm of host Toll signaling-dependent tumor remission shown here appears comparable with Toll-mediated tissue surveillance by microbial infection in *Drosophila* (Germani et al., 2018), or tumor remission induced by Coley’s toxins (Coley) or by TLR agonists (Michaelis et al., 2019).

Serpins have evolved and diversified during the course of animal evolution and are seen to regulate a variety of cellular processes implicated in animal development as well as in host defense (Reichhart, 2005); their subversions may thus trigger cancer progression or metastasis (Valiente et al., 2014). It is conceivable therefore that serpins may constitute part of the spectrum of the elusive nodes of host genetic regulation of cancer progression. We note that forward genetic approaches in the mouse model to dissect TLR-signaling pathways (Beutler et al., 2006) have now identified multiple crosstalk between innate immunity and cancer resistance (Fels Elliott et al., 2017) or susceptibilioty (El-Omar et al., 2008). Our leads from *Drosophila* model—revealing a novel paradigm of host-genetic tumor remission (see Graphical summary)—may further help unearth these elusive genetic factors (Klein, 2009), including serpins, which dictate the fate of an incipient tumor.

## Author Contributions

RS, SH, PS designed the study. SH revealed selective gain of Toll signaling in *lgl* tumors, RS examined the mechanism and established tumor remission. AB analyzed the array data, SP carried out the transcriptomic studies, RKP displayed Lgl expression in immune organs. PS wrote the manuscript and all authors read and contributed.

## Acknowledgement

Authors would like to thank Chris Doe and Satoshi Goto, respectively, for Lgl and Spz antibodies. Research in PS lab was supported by Science & Engineering Research Board (SERB), Department of Science and Technology (New Delhi) research grant NO. CO/S5/AB-03/2017. AB was supported by DBT Wellcome Trust India Alliance (IA/E/13/1/501271).

## SUPPLEMENTARY FIGURES

**Figure S1.**
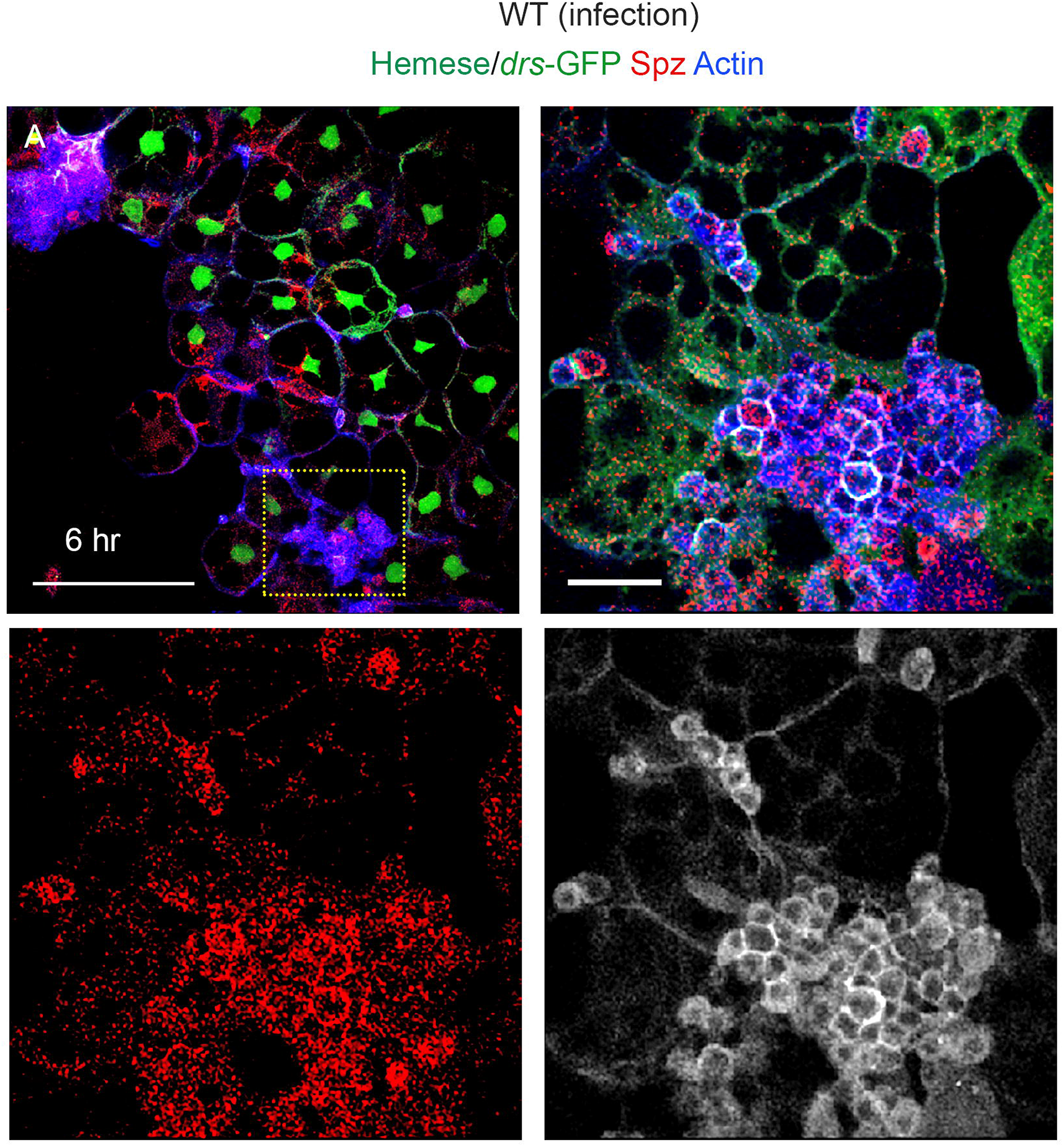
During innate immunity propagation of Toll signaling in larval fat body is mediated by larval hemocytes. **(A)** Fat body of wild type larva upon infection with fungus, *Beauveria bassiana,* displays induction of Toll signaling (*drs-GFP*) in its fat body cells (6 hr after infection, green). Boxed area in (A) is shown at a higher magnification in the rest of the three panels to display that the adherent larval hemocytes (purple, α-Hemese, blue) express the Toll ligand Spz (α-Spz, red). Scale bar= 100μm

**Figure S2.**
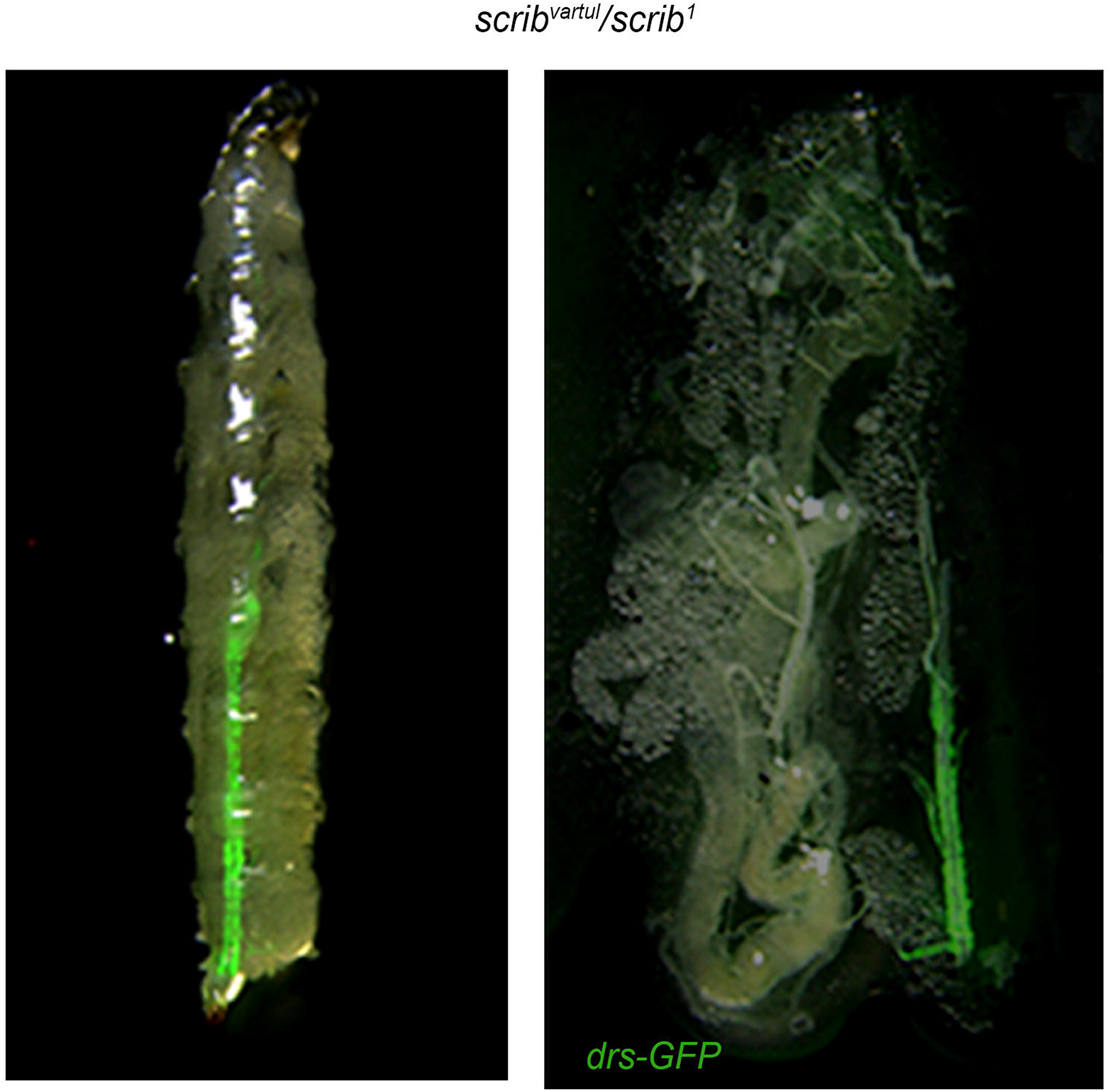
*scrib* mutant larva does not display sterile Toll signaling. *drs-GFP* reporter is not seen *scrib* larval fat body sans its sporadic expression major tracheal branches (n=33/100), reminiscent of that seen following localized loss of a serine protease inhibitor, Spn77BA (Tang et.al, 2008).

**Figure S3.**
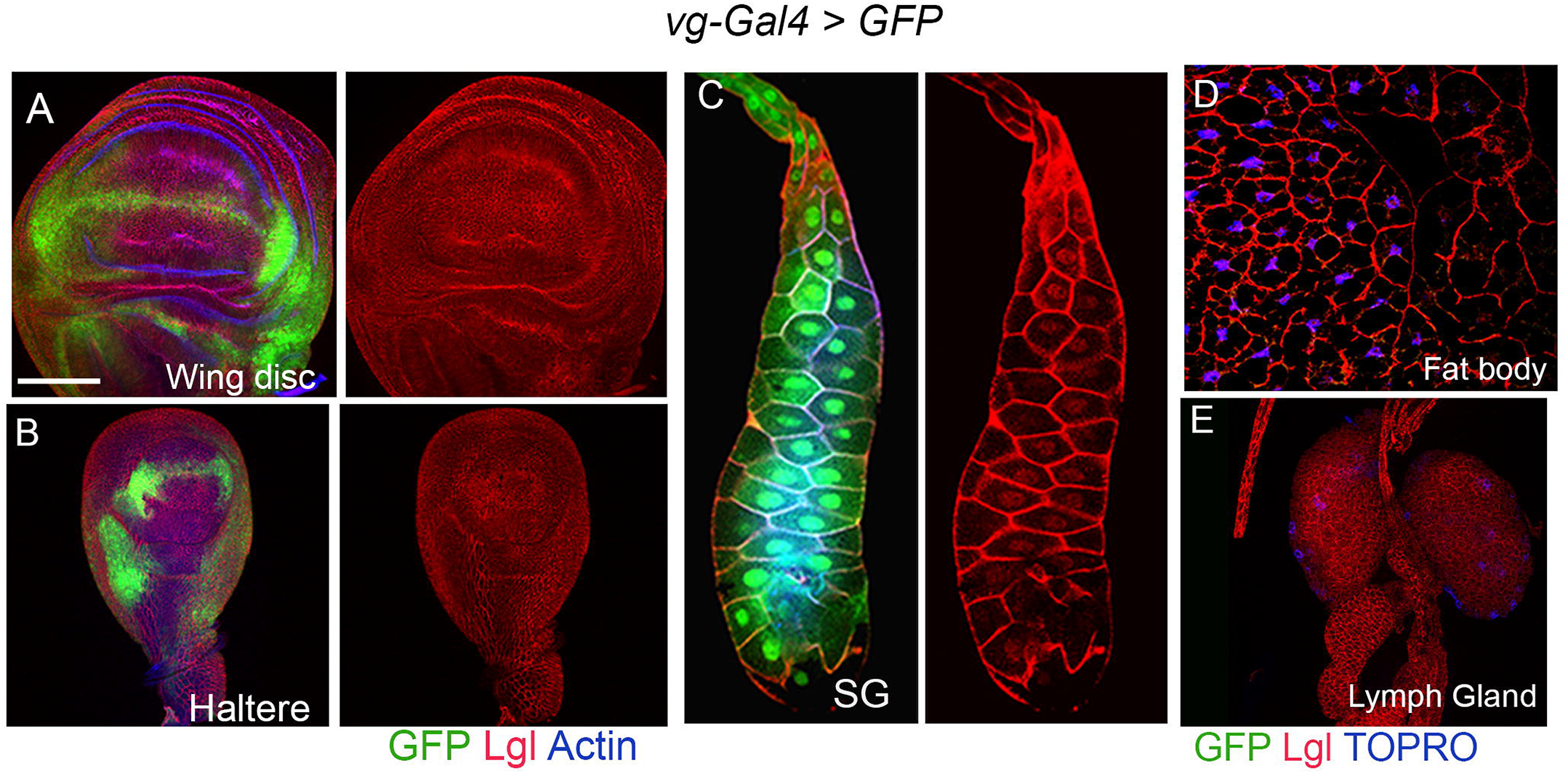
Spatial domains of expression of *vg-Gal4* driver. **(A-E)** Domains of expression *vg-Gal4* driver visualized by a GFP reporter in (A) wing and (B) haltere imaginal discs and in (**C**) larval salivary gland. Note that larval fat body (D) and lymph glands (E) do not express this reporter, while Lgl (red) is expressed in all these organs. Scale bar= 100μm

**Figure S4.**
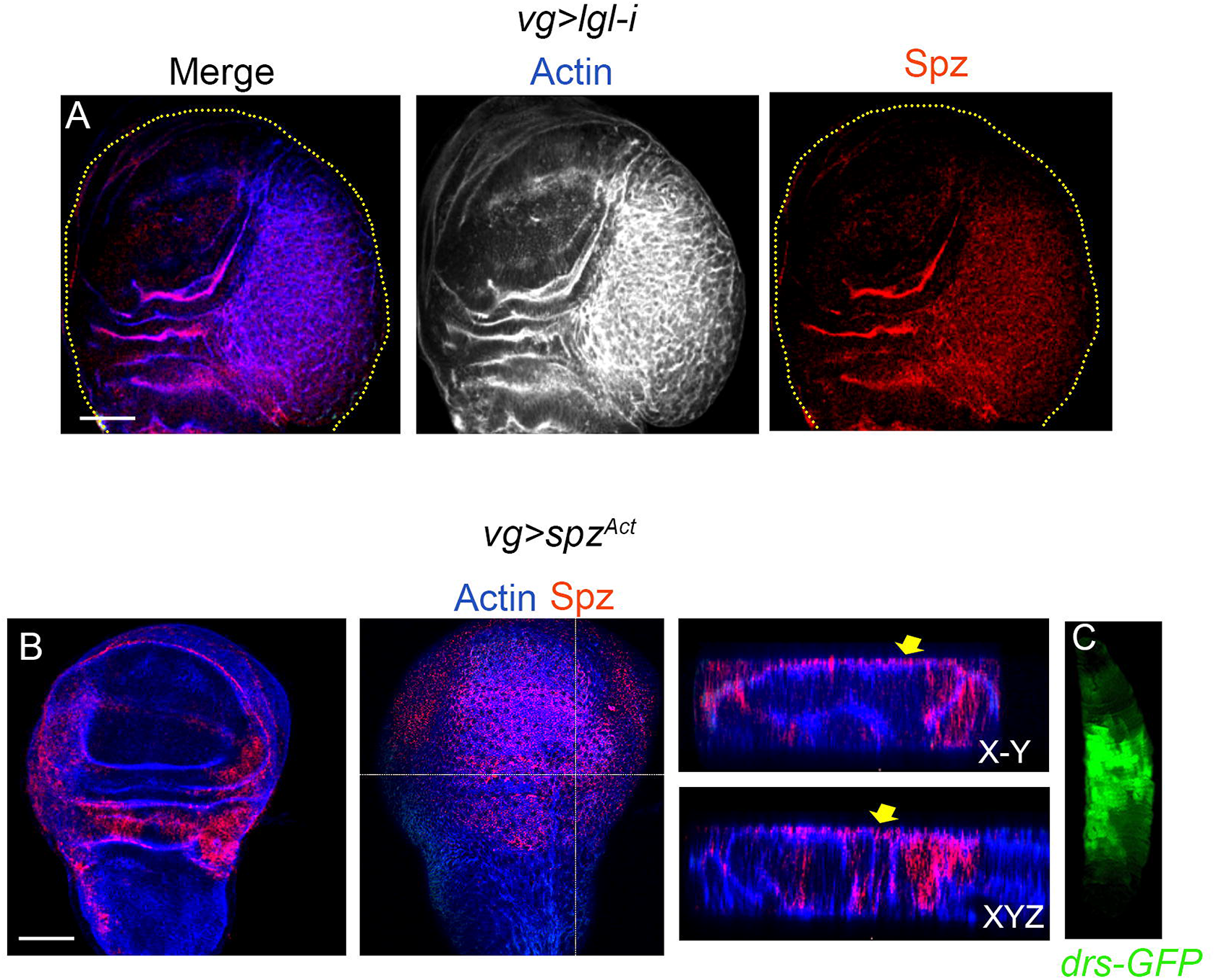
Epithelially produced Spz^Act^ induces systemic Toll signaling. (**A**) Knockdown of *lgl* in wing epithelium under the *vg-Gal4* driver induce neoplastic transformation (star, actin), which also express Spz (red) (**B**) Expression of an activated Spz^Act^ transgene (red) under the *vg-Gal4* driver in wing imaginal disc revealed its secretion of the ligand into the peripodial cells (Wu et al., 2015); X-Y and XYZ along the broken lines are displayed in the right panel to reveal Spz^Act^ (arrow). (**C**) Systemic Toll signaling (*drs-GFP* in fat body) in a wild type larva displaying Spz^Act^ expression under the *vg-Gal4* driver. Scale bar= 100μm

**Figure S5.**
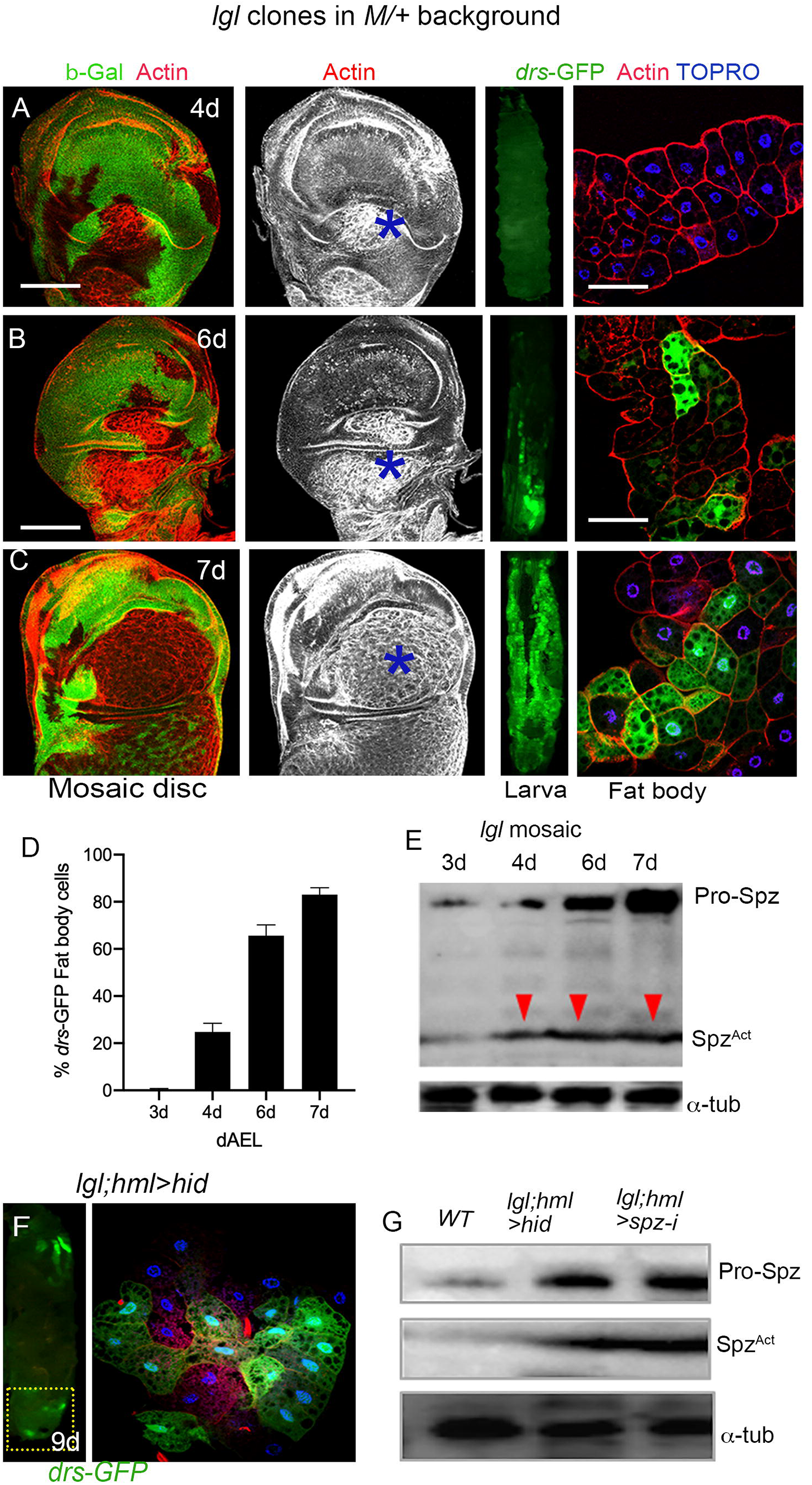
Tumor progression-linked systemic Toll signaling is propagated by host hemocytes. **(A-C).** Somatic clones of *lgl* (absence of GFP) induced in a *Minute/+* genetic background (GFP, green). With progression of larval age, these *lgl* clones displayed neoplastic transformation (actin, star), while fat of the host larvae displayed progressive increase in *drs-GFP* expressing adipocytes in their fat body. (**D**) Quantification adipocytes expressing drs-GFP. (**E**) Western blots of these mosaic imaginal discs to reveal progressive increase in the levels of Spz^Act^ with increasing larval age (tumor progression). (**F**) Ablation of hemocytes (*hml>hid*) in *lgl* larvae arrested propagation of Toll signaling in their fat body (see magnified box area on the right) although their neoplasms (**G**) produced Spz^Act^. Larval age is shown in <u>d</u>ays <u>a</u>fter <u>e</u>gg <u>l</u>aying (dAEL). Scale bar=100μm.

**Figure S6.**
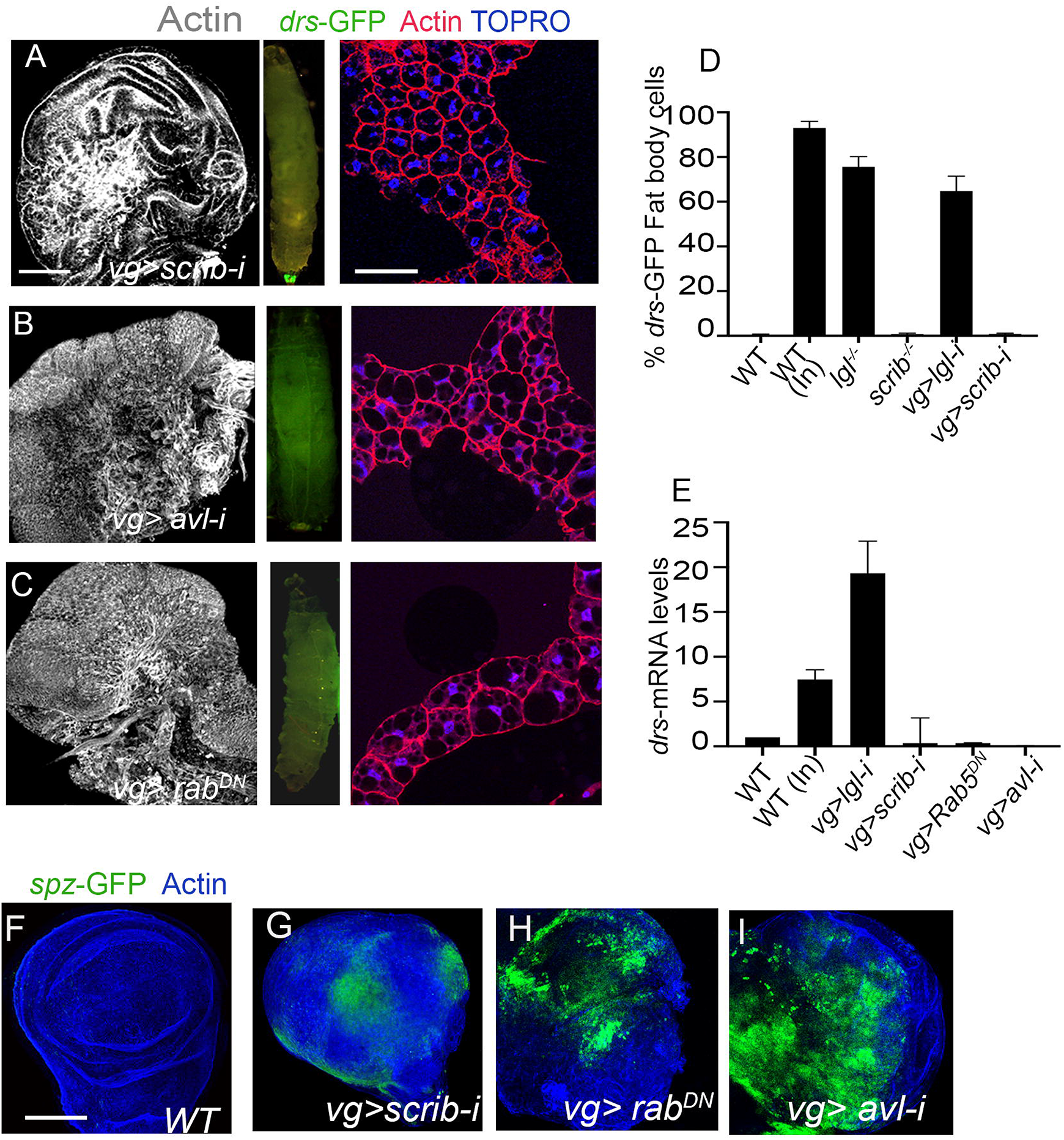
Select induction of Systemic Toll signaling by *lgl* tumors. (**A-C)** Knockdown of *scrib* (A), *avl* (B) expression of a dominant negative *Rab5*^*DN*^ transgene (C) under the *vg-Gal4* driver induce extensive neoplastic transformation of wing epithelium. The host larval fat body, however, do not display expression of the *drs-GFP* reporter (D) Quantification of *drs*-*GFP* reporter and (E) *drs* m-RNA in fat body of relevant genotypes following infection (In) or when raised in axenic medium. (**F**-**I)** Comparison *spz-GFP* reporter expression in wild type wing epithelium (F) with those shown in (A-C). Scale bar= 100μm

**Figure S7.**
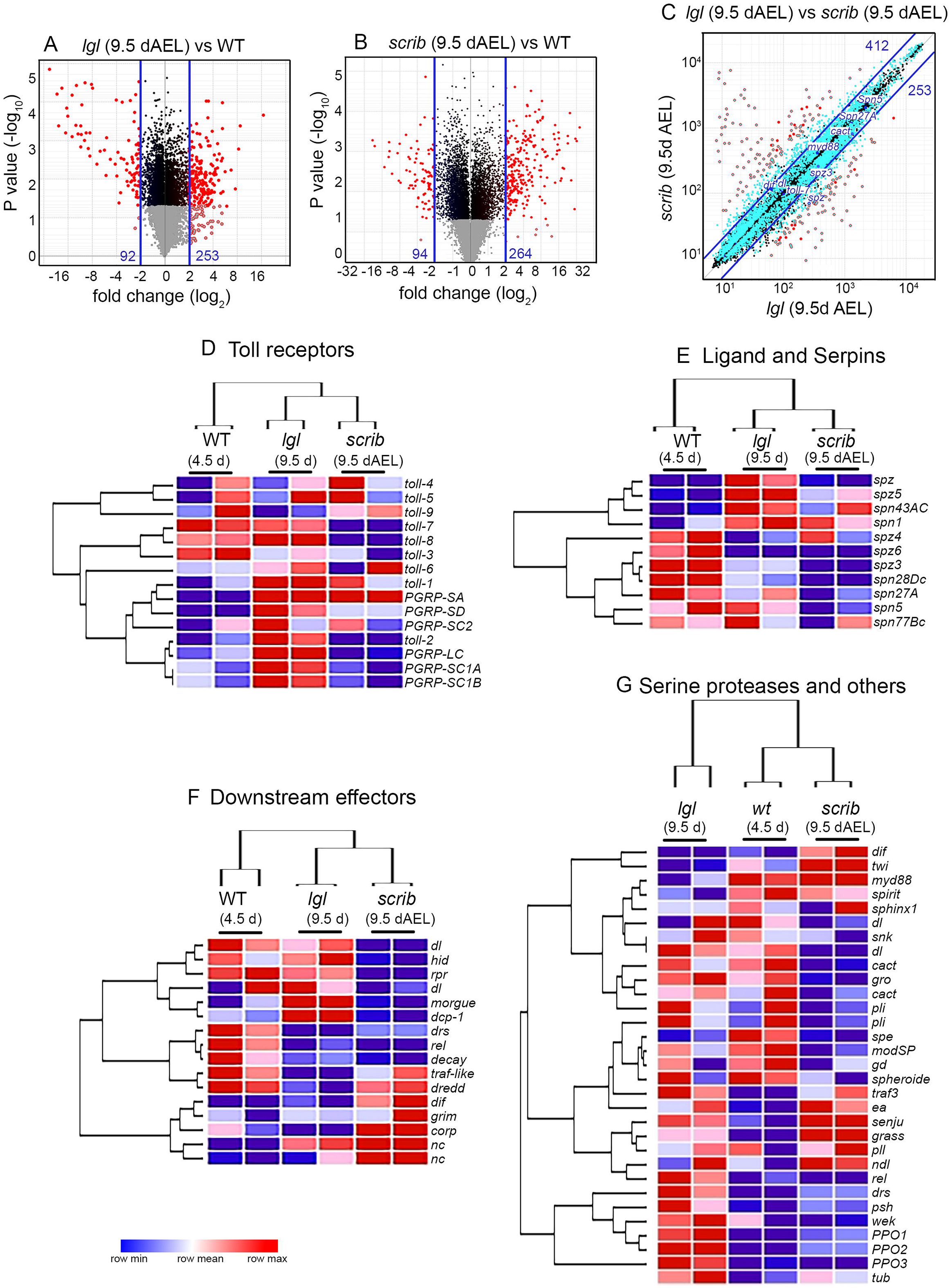
Transcriptional statuses of Toll signaling pathway members in transcriptomes of *lgl* and *scrib* neoplasms. (A-B) Volcano plot depicting fold change in gene expression in *lgl* (A), and *scribble* (B) tumors at 9.5 days, when compared to their wild type (4.5 day) control discs. Each point represents the average fold change in expression for a single gene (log_2_, x-axis) at a P value (−log_10_, y axis) in tumor versus wild type (WT) controls. Genes with significant fold change in expression (>± log_2_ 2-fold change, P<0.05) are marked in red (outside the blue solid line), their total number denoted on either side of the blue lines. (C) Correlation plot comparing expression of genes in *lgl* (x axis) and *scrib* (y axis) tumors of comparable age (9.5 days). Blue solid lines along the diagonal demarcate genes with expression differences of more than 2-fold, while data points in red mark genes with ± 4-fold difference in gene expression between *lgl* and *scrib* tumors. Data points representing some of the Toll pathway members are highlighted; note that these largely lie within the blue lines which point to moderate difference in their expression between the *lgl* and *scrib* tumors. (**D-G**) Hierarchical clustered-heatmaps with row normalization displaying transcriptional statuses of genes coding for Toll signaling pathway members: namely, Toll receptors (D), Toll ligands and Serpins (E) and downstream effectors (F) besides other Toll-associated members including SPs (G) in 9.5 dAEL neoplasms of *lgl* and *scrib* mutants and 4.5 dAEL WT wing discs. Note the upregulation of Toll receptors like *toll-2, toll-1* and *toll-6* (D), and Toll ligands, Spz and Spz5 (E), in *lgl* neoplasms compared to the control WT discs. SP processing enzymes, Serpins, were either upregulated (*spn1, spn43C*) or downregulated (*spn28Dc*) in both *lgl* and *scrib* tumors (E), compared to WT discs, while SPs such as *ea*, *psh*, *grass*, *pll* were upregulated in both (G), although Mod-SP was selectively downregulated in *scrib* neoplasm (G). Effectors of Toll pathway, NFkB homolog *dl* was upregulated in *lgl* whereas *dif* was upregulated in *scrib* neoplasms as compared to WT discs (G). Cell death-associated genes *Dcp-1* and *morgue,* including proapoptotic genes *hid*, *reaper* was downregulated in *scrib* tumors, while these were upregulated in *lgl* tumors (F).

**Figure S8.**
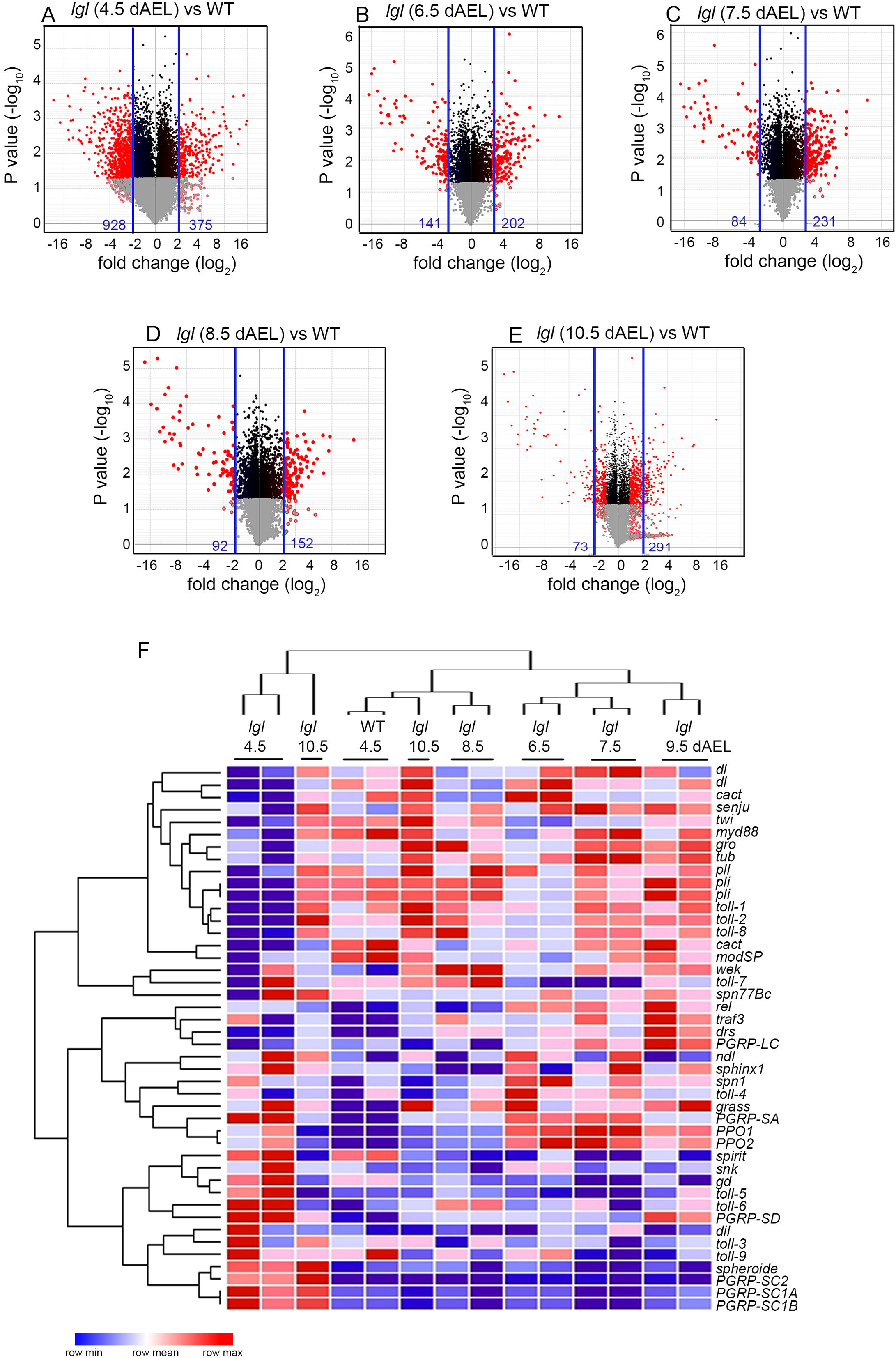
Transcriptional statuses of members of Toll signaling pathway during *lgl* tumor progression. (**A-E**) Volcano plots representing differential gene expression during *lgl* progression (4.5 to 105 dAEL) versus wild type (WT) third instar wing imaginal discs. Each point represents the average fold change in expression for a single gene (log_2_, x-axis) at a P value (−log_10_, y axis) in tumor versus WT control. Genes with significant fold changes in expression (>± log_2_ 2-fold change, P<0.05) are marked in red (outside the blue solid line), their total numbers are denoted on either side of the blue lines. (F) Hierarchically clustered gene expression heatmaps of members of Toll signaling pathway in progressively aged (4.5 to 10.5 dAEL) *lgl* neoplasms compared to those of WT wing imaginal disc on 4.5 dAEL. Overall, expression levels of Toll pathway members in early stage of *lgl* tumor progression (4.5 dAEL, columns 1 and 2) were strikingly different from those of at later time points. We noted two distinct clusters of genes: those that were transiently down-regulated (top rows in the heat map) and those that were up-regulated (bottom rows in the heat map) in 4.5day *lgl* neoplasms. For instance, genes coding for Toll receptors (*toll-1, toll-2* and *toll-8),* effectors, or negative regulators of the NF-κB (*cact*), displayed a transient (4.5 dAEL) down-regulation while those coding for *toll-5*, *toll-6* and *PGRPs* were transiently up-regulated in 4.5D AEL neoplasms.

**Figure S9.**
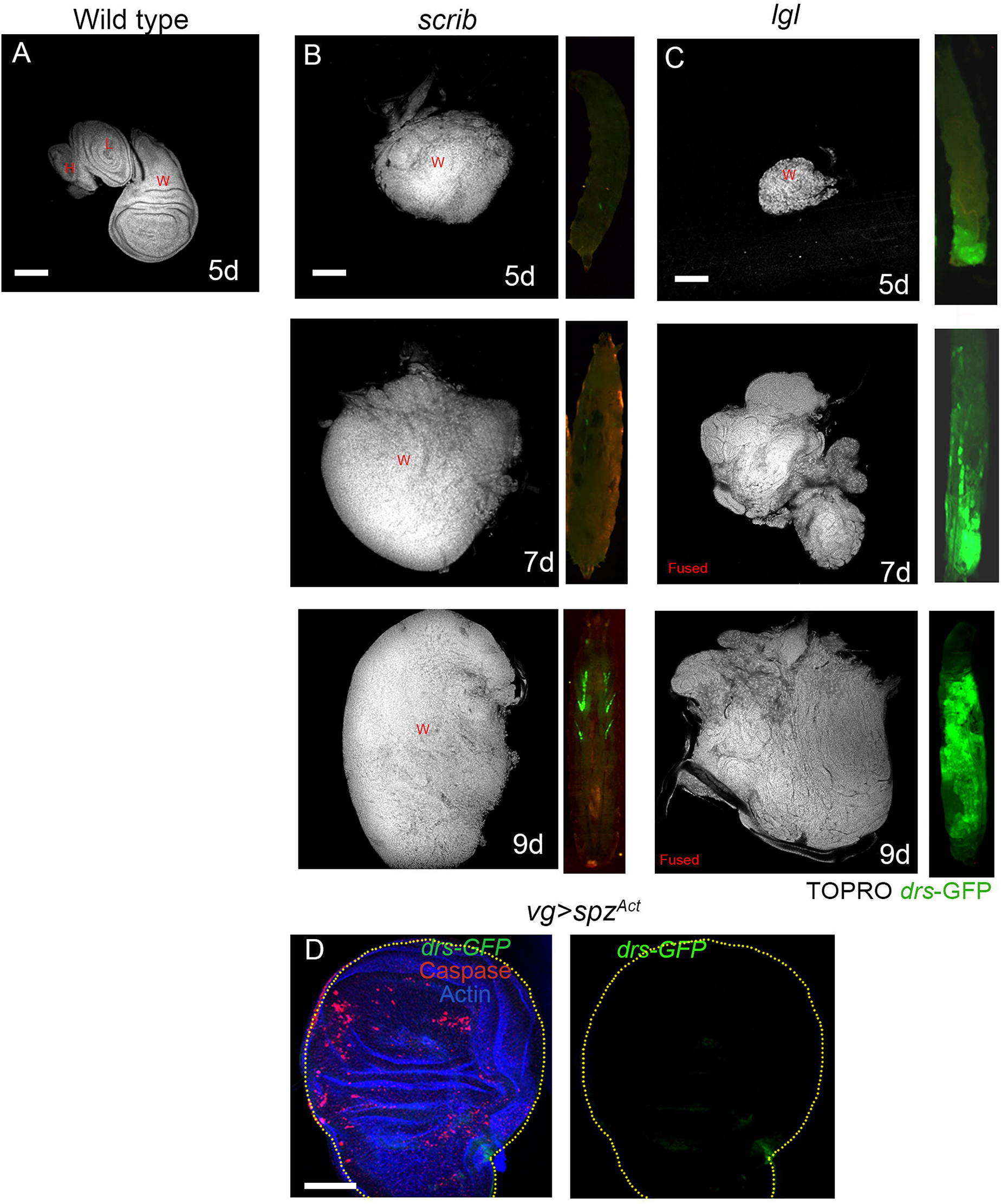
*lgl* neoplasm display staggered tumor progression as compared to their *scrib* counterparts. (**A- C**) Wild type wing, haltere and leg imaginal discs (A) as compared to neoplasms of scrib (B) and *lgl* (C) wing imaginal disc on 5 dAEL During subsequent days of tumor progression (7-9 dAEL), neoplasms of *scrib* mutant wing imaginal disc display massive increase in the sizes (B), while those of *lgl* display fusion/invasion into each other that obliterates their individual identity (C). (**D**) Expression of a *spz*^*Act*^ transgene under the *vg-Gal4* driver induced cell death (caspase) in wild type wing imaginal disc, while Toll innate immunity reporter, drs*-GFP,* remains silent. Abbreviations: W=wing, H=haltere and L=leg. Larval age is displayed as dAEL. Scale bar= 100μm

**Figure S10.**
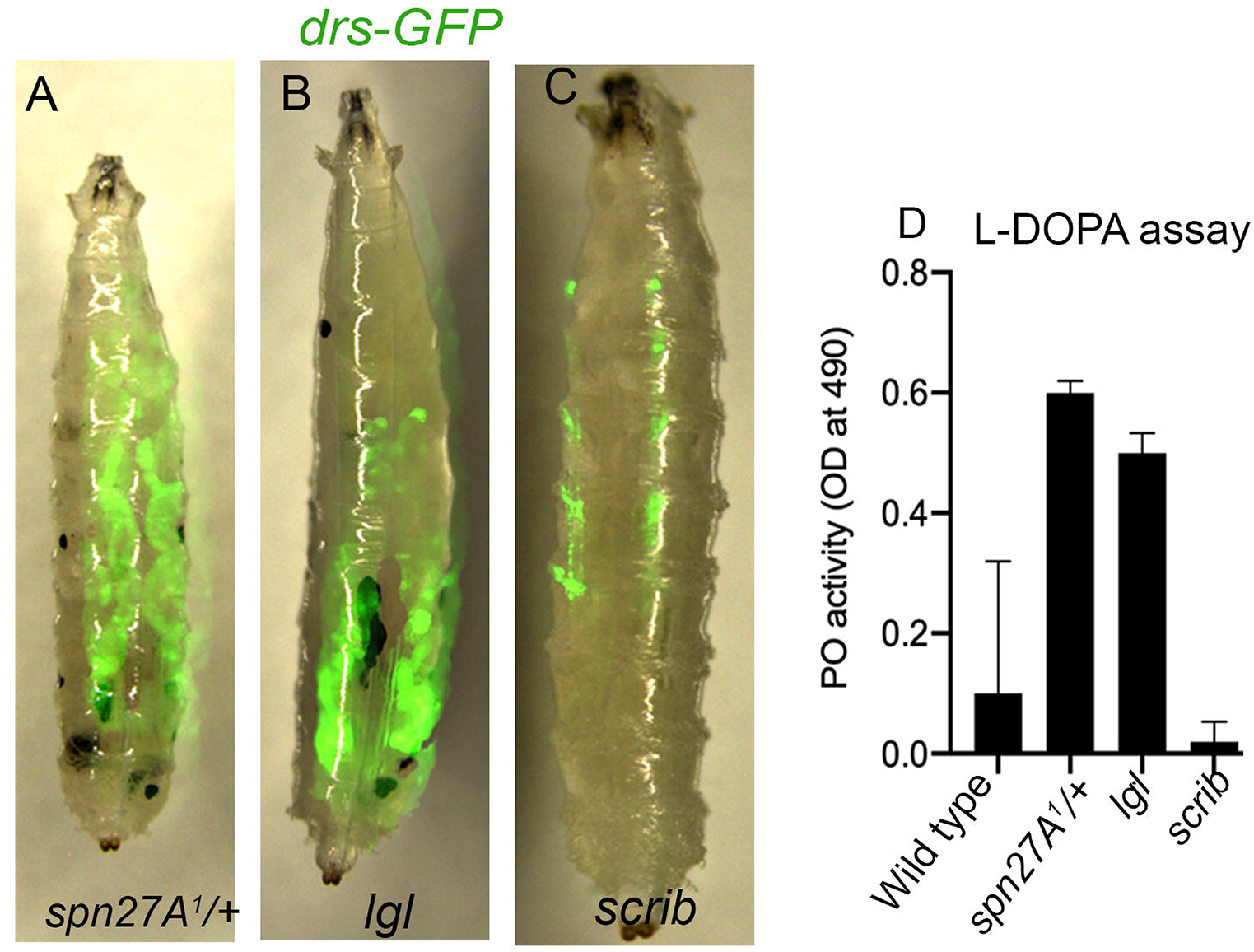
*spn27A*^*1*^/*+* host larvae and adults display constitutive Toll signaling. (**A-D**) Both *spn27A*^*1*^/*+* (A) and *lgl* (B) larvae display spontaneous (constitutive) melanization in their hemolymph marked by excessive production of melanin due to high level of an activated serine protease, phenol oxidase (PO) (Ligoxygakis et al., 2002), besides activation of the *drs-GFP* expression in the larval fat body; their *scrib* counterparts do not display these hall marks of *spn27A* down regulation or activation of SPs (C). L-DOPA assay in hemolymph of these larvae provide a quantification of their respective PO activity (D). Scale bar= 100μm.

**Figure S11.**
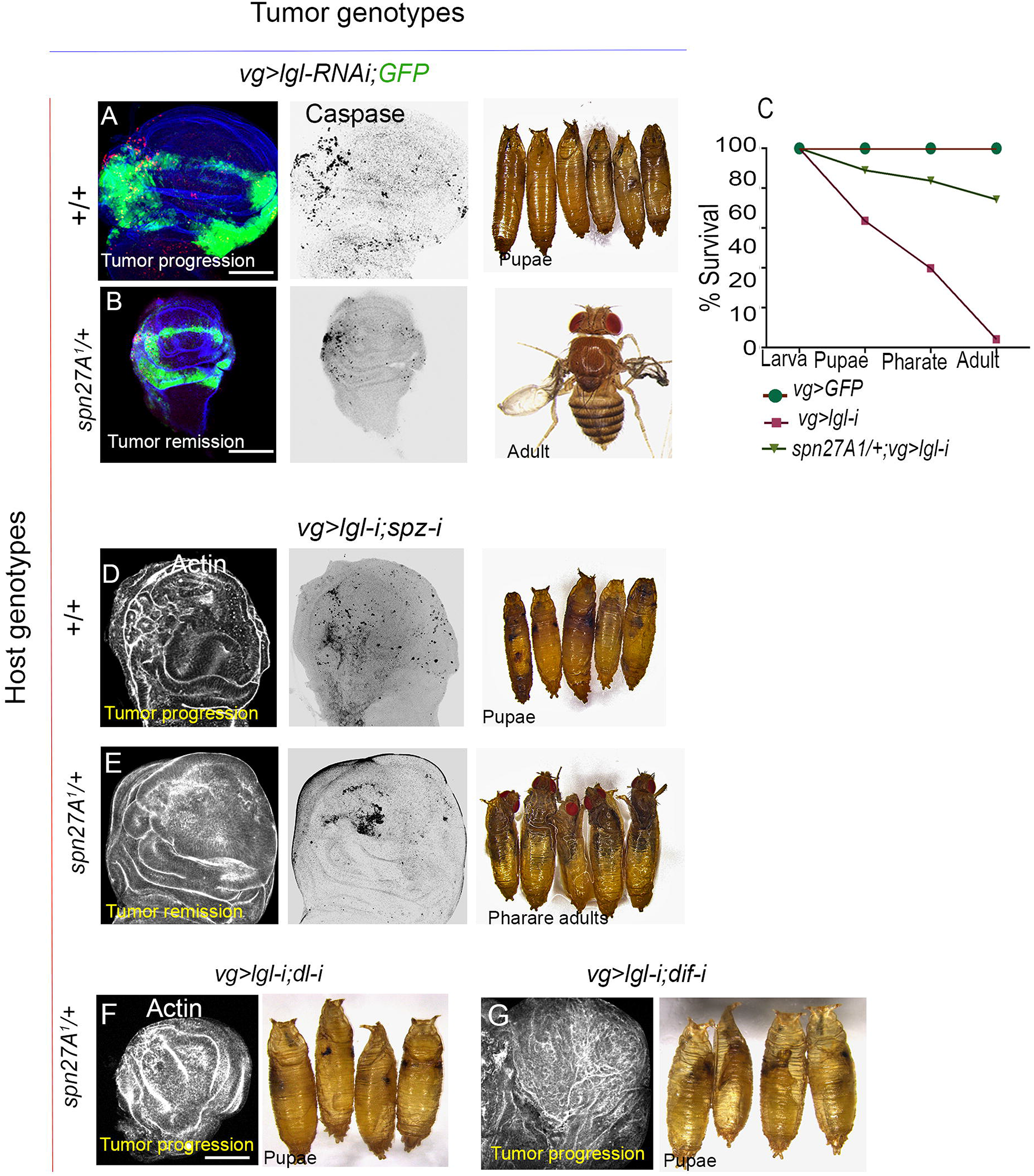
Toll-mediated remission of *lgl* neoplasms in *spn27A*^*1*^/*+* host genetic background. (**A**-**E**) *lgl* wing imaginal disc neoplasms (actin) display rare cell death (caspase) in wild type host larvae, which die as pupae (A). Conversely, *lgl* neoplasms in s*pn27A*^*1*^/*+* larval hosts display extensive cell death (caspase), tumor remission and survive as adults, which display characteristic Toll signaling (d*rs-GFP*) survival plots (C) of the animals shown in (A, B). *lgl* neoplasms displaying knockdown of *spz* displayed rare cell death (caspase) in wild type host larvae, which died as pupae (D), while these neoplasms are eliminated by cell death (caspase) in s*pn27A*^*1*^/*+*larval hosts that survived till pharate adult stage (E). (**F, G**) Restoration of tumor progression upon blocking reception of Toll signaling (*dl-i,* F or *dif-i,*G) in *lgl* neoplasms induced in s*pn27A*^*1*^/*+* larval hosts, which then displayed pupal lethality. n (5) =100 for each set. Scale bar=100μm.

**Figure S12.**
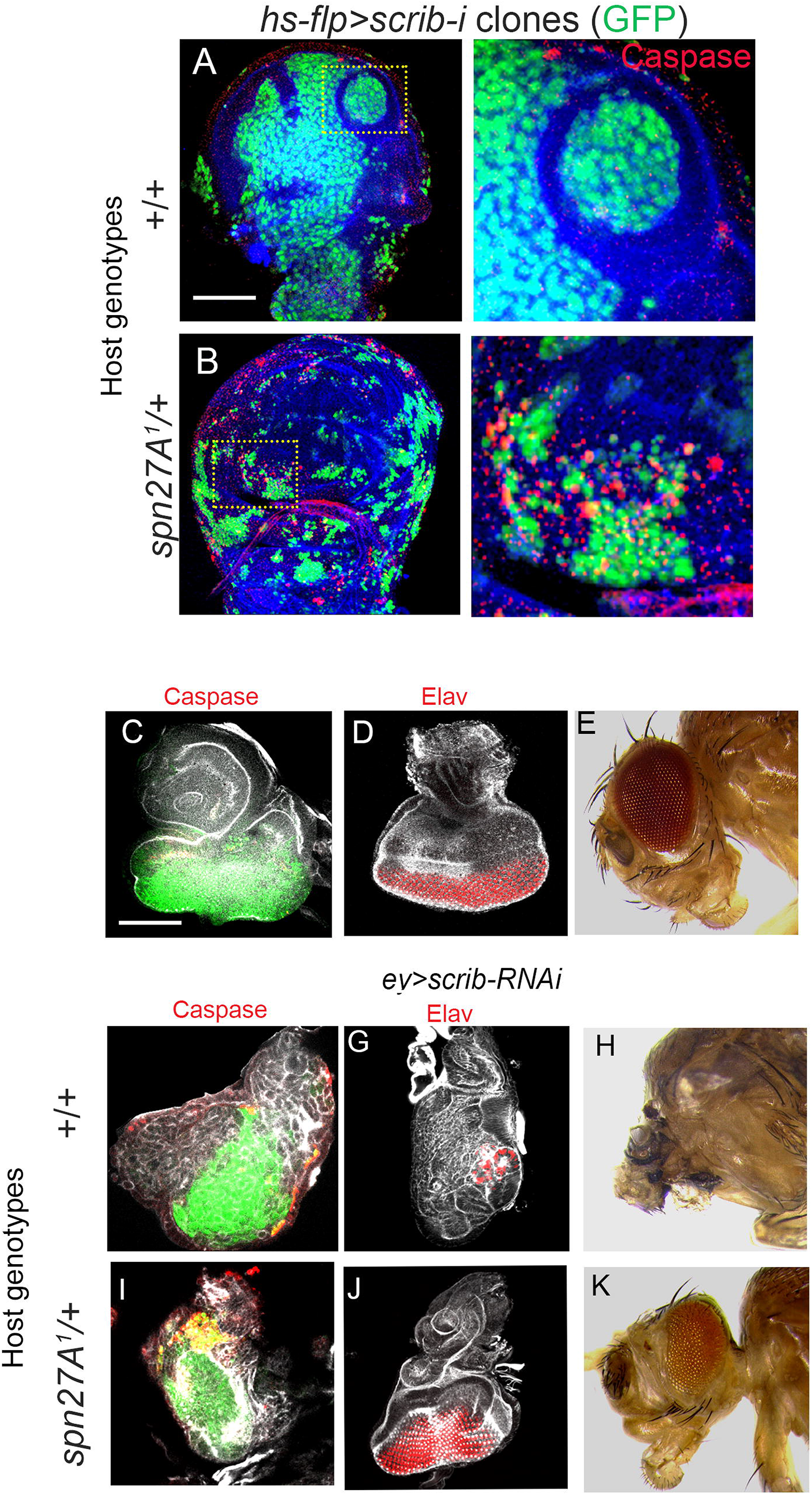
Host genetic Toll signaling induces remission of multiple types of *Drosophila* neoplasms. (**A, B**) *scrib* clones generated in wing imaginal discs in wild type (*+/+*, A) or in *spn27A*^*1*^/*+* (B) host genetic background. Note cell death (caspase, red) in *scrib* clones (green) induced in the *spn27A*^*1*^/*+* hosts. (**C**-**E**) Wild type eye-antennal discs stained for caspase (red), actin (grey) and, ommatidial cell differentiation (Elav, red, D). Wild type adult head (E). (**F-K**) *scrib* neoplasms (*ey>scrib-i)* induced in wild type(F-H) and in *spn27A*^*1*^/*+* (I-K) genetic background (n=220/300 for adult eclosion). Scale bar= 100μm

## Material and methods

### Fly husbandry

Fly stocks were grown on standard axenic corn meal-agar medium supplemented with streptomycin (400μg/ml) along with anti-fungal Antimycotic Nipagin (0.15 g per 100 ml), at 25 °C including those involving Gal4 driver lines and instances of shift to 18± 1°C at 4.5 dAEL (days <u>A</u>fter <u>E</u>gg <u>L</u>aying) are mentioned wherever done. Care has taken to raise larvae and flies under aseptic and uncrowded conditions, especially when assaying for systemic immune responses (also, see Germani et al., 2018).

### *Drosophila* stocks

Fly strains used in present study including mutant lines, reporter lines carrying GFP marked chromosomes, Gal4 drivers, UAS- and RNAi lines were received from Bloomington *Drosophila* Stock Centre (BDSC) and Vienna *Drosophila* Resource Center (VDRC), while some were received as gifts from other investigators.

### Genotypes

For assessing *drosomycin* expression, a *drs-GFP* reporter was introgressed in multiple genetic backgrounds. Genotypes of stocks used are listed below:

**Figure 1**

A, B: **Control/wild type**= *w*^*118*^; *drs-GFP*

C, D: ***hml>spz-i***= *w*^*118*^; *hml-Gal4>UAS-dicer RNAi; UAS-spz RNAi; drs-GFP*

E: ***lgl***= *w*^*118*^; *lgl*^*4*^; *drs-GFP*

F: ***scrib***= *w*^*118*^; *scrib*^*vartul*^/*scrib*^*1*^; *drs-GFP*

G, I: ***vg>lgl-i***= *w*^*118*^; *vg-GAL4>UAS-dicer RNAi; UAS-lgl RNAi, UAS-GFP*

H, J: ***vg>scrib-i***= *w*^*118*^; *vg-GAL4>UAS-dicer-RNAi; UAS-scrib-RNAi, drs-GFP*

K: ***lgl; hml>spz-i***= *w*^*118*^; *lgl*^*4*^, *hml-Gal4>UAS-spz-RNAi, drs-GFP*

L: ***vg>lgl-i; spz-i***= *w*^*118*^; *vg-GAL4>UAS-dicer-RNAi; UAS-lgl-RNAi, UAS-spz-RNAi, drs-GFP*

**Figure 2**

D: ***vg>lgl-i; spz-i*** = *w*^*118*^; *vg-GAL4>UAS-dicer-RNAi; UAS-lgl-RNAi, UAS-spz-RNAi, drs-GFP*

E: ***vg>lgl-i; dl-i***= *w*^*118*^; *vg-GAL4>UAS-GFP, UAS-dicer-RNAi; UAS-lgl-RNAi, UAS-dl-RNAi*

G: ***vg>lgl-i; spn27A***= *w*^*118*^; *vg-GAL4>UAS-spn27A; UAS-lgl-RNAi, drs-GFP*

H: ***vg>scrib-i; spn27A-i***= *w*^*118*^; *vg-GAL4>UAS-spn27A-RNAi; UAS-scrib-RNAi, drs-GFP*

**Figure 3**

A: ***vg>scrib-i***

Genotype of neoplasm: *vg-GAL4>UAS-scrib-RNAi*

Genotype of Host larval: *w*^*118*^ (*+/+*), *drs-GFP*

B: ***spn27A***^***1***^/+; ***vg>scrib-i***

Genotype of the neoplasm: *vg-GAL4>UAS-scrib-RNAi*

Host larval genotype: *spn27A*^*1*^/+; *drs-GFP*

D: ***vg>scrib-i; spz-i***

Genotype of the neoplasm: *vg-GAL4>UAS-scrib-RNAi, UAS-spz-RNAi*

Genotype of Host larval: *w*^*118*^ (*+/+*), *drs-GFP*

E: ***spn27A***^***1***^/+; ***vg>scrib-i; spz-i***

Genotype of the neoplasm: *vg-GAL4>UAS-scrib-RNAi, UAS-spz-RNAi*

Genotype of Host larval: *spn27A*^*1*^/+, *drs-GFP*

F: ***spn27A***^***1***^/+; ***vg>scrib-i; dl-i***

Genotype of the neoplasm: *vg-GAL4>UAS-scrib-RNAi, UAS-dl-RNAi*

Genotype of Host larval: *spn27A*^*1*^/+, *drs-GFP*

G: ***spn27A***^***1***^/+; ***vg>scrib-i; dif-i***

Genotype of the neoplasm: *vg-GAL4>UAS-scrib-RNAi, UAS-dif-RNAi*

Genotype of Host larval: *spn27A*^*1*^/+, *drs-GFP*

**Figure S1:**

A: **Control/wild type**= *w*^*118*^; *drs-GFP*

**Figure S2:**

A: ***scrib***= *w*^*118*^; *scrib*^*vartul*^/*scrib*^1^; *drs-GFP*

**Figure S3:**

A: ***vg>GFP/wild type*** = *w*^−^; *vg-GAL4>UAS-GFP*

**Figure S4:**

A, B: ***vg>lgl-i*** = *w*^*118*^; *vg-GAL4>UAS-dicer-RNAi; UAS-lgl-RNAi*, *drs-GFP*

C, D: ***vg>spz*^*Act*^** = *w*^*118*^; *vg-GAL4>UAS-spz*^*Act*^; *drs-GFP*

**Figure S5:**

A-E: ***lgl***^−^ ***M***^+^ : *y w hs-flp; lgl*^*4*^ *FRT40,Minute arm-lacZ* FRT40 *; drs*-GFP

F: ***lgl;hml>hid***= *w*^*118*^; *lgl*^*4*^,*hml-Gal4> UAS-hid*, *drs-GFP*

**Figure S6:**

A: ***vg>scrib-i*** = *w*^*118*^; *vg-GAL4>UAS-dicer-RNAi; UAS-scrib-RNAi*, *drs-GFP*

B: ***vg>avl-i*** = *w*^*118*^; *vg-GAL4>UAS-dicer-RNAi; UAS-avl-RNAi*, *drs-GFP*

C: ***vg>Rab5*^*DN*^** = *w*^*118*^; *vg-GAL4>UAS-Rab5*^*DN*^, *drs-GFP*

F: **wild type**= *w*^*118*^; *spz-GFP*

G: ***vg>scrib-i*** = *w*^*118*^; *spz-GFP, vg-GAL4>UAS-dicer-RNAi; UAS-scrib-RNAi*

B: ***vg>avl-i*** = *w*^*118*^; *spz-GFP, vg-GAL4>UAS-dicer-RNAi; UAS-avl-RNAi*

C: ***vg>Rab5*^*DN*^** = *w*^*118*^; *spz-GFP, vg-GAL4>UAS-Rab5*^*DN*^

**Figure S9:**

A: **Control/wild type**= *w*^*118*^; *drs-GFP*

B: ***scrib***= *w*^*118*^; *scrib*^*vartul*^, *scrib*^*1*^; *drs-GFP,*

C: ***lgl***= *w*^−^; *lgl*^*4*^; *drs-GFP*

**Figure S11:**

A: ***vg>lgl-i***

Genotype of the neoplasm: *vg-GAL4>UAS-lgl-RNAi*

Genotype of Host larval: *w*^*118*^ (*+/+*), *drs-GFP*

B: ***spn27A***^*1*^/+; ***vg***>***lgl-i***

Genotype of the neoplasm: *vg-GAL4>UAS-lgl-RNAi*

Genotype of Host larval: *spn27A*^*1*^/+, *drs-GFP*

C: ***vg>lgl-i; spz-i***

Genotype of the neoplasm: *vg-GAL4>UAS-lgl-RNAi, UAS-spz-RNAi*

Genotype of Host larval: *w*^*118*^ (*+/+*), *drs-GFP*

D: ***spn27A***^***1***^/+; ***vg>lgl-i; spz-i***

Genotype of the neoplasm: *vg-GAL4>UAS-lgl-RNAi, UAS-spz-RNAi*

Genotype of Host larval: *spn27A*^*1*^/+, *drs-GFP*

E: ***spn27A***^***1***^/+; ***vg>lgl-i; dl-i***

Genotype of the neoplasm: *vg-GAL4,>UAS-lgl-RNAi, UAS-dl-RNAi*

Genotype of Host larval: *spn27A*^*1*^/+, *drs-GFP*

F: ***spn27A***^***1***^/+; ***vg>lgl-i; dif-i***

Genotype of the neoplasm: *vg-GAL4>UAS-lgl-RNAi, UAS-dif-RNAi*

Genotype of Host larval: *spn27A*^*1*^/+, *drs-GFP*

**Figure S12:**

A: ***hs-flp>scrib-i***

Genotype of the neoplasm: *hs-flp>UAS-scrib-RNAi, UAS-GFP*

Genotype of Host larval: *w*^*118*^ (*+/+*), *drs-GFP*

B: ***spn27A***^***1***^/+; ***hs-flp>scrib-i***

Genotype of the neoplasm: *hs-flp>UAS-scrib-RNAi, UAS-GFP*

Genotype of Host larval: *spn27A*^*1*^/+, *drs-GFP*

C: ***ey>GFP/wild type***= *w*^*118*^; *ey-GAL4>UAS-GFP*

D: ***ey>scrib-i***

Genotype of the neoplasm: *ey-GAL4>UAS-scrib-RNAi, UAS-GFP*

Genotype of Host larval: *w*^*118*^ (*+/+*), *drs-GFP*

E: ***spn27A***^*1*^/+; ***ey>scrib-i***

Genotype of the neoplasm: *ey-GAL4>UAS-scrib-RNAi, UAS-GFP*

Genotype of Host larval: *spn27A*^*1*^/+, *drs-GFP*

### Fungal strains and Culture

*Beauveria bassiana*, an entomopathogenic fungus, that is known to act as a parasite on various arthropods, including *Drosophila* procured from Institute of Microbial Technology, Chandigarh. Standard culture of *Beauveria bassiana* (BbM1), strain MTCC-2028 was grown on potato dextrose medium at 37°C.

### Larval infection

Third instar larvae grown on axenic culture conditions were harvested and washed three times sterile distilled water and one time in Phosphate Buffer Saline (PBS, pH-7.4). These were then placed in an ice-cold glass plate to reduce their movement, and fungal mycelium of *Beauveria bassiana* was injected using a sterile tungsten needle through the cuticle between larval body segments 7 and 8, which were then axenic medium for further study.

### Whole larval imaging

Larvae carrying *drs-GFP* reporter were washed with Phosphate Buffer Saline (PBS, pH 7.4) and immobilized (kept on ice) and viewed under epi-fluorescent illumination (excitation filter 480 nm; dichroic filter 505 nm LP; emission filter 510 nm LP) with a Leica M205FA stereomicroscope set up with digital camera Leica DC300 F. Larval fat bodies were dissected in PBS *drs*-GFP fluorescence was captured.

### Generation of mutant clones in wing imaginal disc epithelium

Somatic clones were generated by *flp*-FRT and MARCM techniques (Khan et al., 2013). Heat shock was given 2dAEL (days <u>A</u>fter <u>E</u>gg <u>L</u>aying) at 37°C for 30 minutes to induce *hs-flp* mediated somatic recombination. These larvae were dissected at different time-points, which varied from 3d to 7 dAEL.

### Larval imaginal disc and fat body dissection and immunostaining

Wandering third instar larvae were picked for imaginal disc/fat body dissection and first thoroughly washed in Phosphate Buffered Saline (PBS) to get rid of food particle and other debris. First, posterior end incision was made in such that the hemolymph and gut tissues were squeezed out using a pair of needles. Next, larval cuticle inside-out, exposing out the imaginal discs, fat body, brain lobes and other organs.

Third instar flipped larvae were transferred to 1ml centrifuge tube and fixed in 4% para-formaldehyde in PBS at 4°C for 1 hours. Fixative was removed by multiple washes with PBS containing 0.2% Triton X-100 (PBST) to permeabilize the membrane to achieve immunostaining. Fixed samples were blocked in PBS containing 10% Bovine Serum Albumin for 2-4 hours before incubating with primary antibodies overnight at 4°C. Primary antibodies used in this study are Anti-Lgl (rabbit, 1:300, gift from Chris Doe), anti- Spz (rat, 1:50, requested from Satoshi Goto), Anti-Elav (rabbit, 1:50, DSHB) and Caspase 3 (rabbit, 1:500, Sigma Aldrich). Samples were washed four times for 10 minutes each in PBST and incubated at room temperature in secondary antibodies: Anti-rat alexa Fluor-555, anti-rabbit Alexa Fluor-555 (purchased from Invitrogen Inc.) for 2 hrs. Washes were given in PBST four times for 10 minutes. Fat body tissue was dissected in PBS, and mounted in Vectashield mounting medium (Vector Labs, H1200).

Fat body tissue/ imaginal disc was dissected, fixed, washed, and blocked in 10% BSA in PBT as described above before incubating in Phalloidin-555, for 35-40 minutes. The sample was then washed four times for 20 minutes in PBST and then incubated in TO-PRO-3, the nuclear labelling dye for 1 hour at room temperature before mounting.

### Immunofluorescence and microscopy

All fluorescence imaging was done with Leica-SP5 Confocal microscope and processed using Leica Confocal Software-LAS AF and Adobe Photoshop.

### *drs-GFP* positive Fat body cell quantification

Fat body images were scored for nuclear *drs-GFP* using LAS-AF confocal software. Bar graphs and statistical significance scores were generated on GraphPad using paired *t* test.

### Tumor size measurement

Tumors were mounted in Vectashield and confocal micrographs were taken at mid-depth of the sample. The area occupied by tumor was measured using ImageJ (NIH).

### Western immunoblotting

Ten larvae were washed first in70% ethanol then twice with sterile PBS at room temperature. The larvae were then pricked with a sterile capillary to collect the hemolymph, which was suspended in 30 μL of lysis buffer (RIPA buffer).

Wing imaginal disc epithelium or their neoplasms, were dissected from 50 larvae of these were dissected in PBS and collected in 50 μL of lysis buffer, and sonicated (3 cycles of the 30 sec pulse with 30 sec interval on ice) using Bioruptor (Diagenode). The mixture was then boiled at 95 °C for 5 min to denature the proteins, and centrifuged at 15,000 g (4 °C) for 15 min to separate the cell debris. The supernatant was collected for further analysis. Approximately ~80-100 μg of the denatured protein isolated from hemolymph or imaginal epithelia/neoplasms from individual genotype were run on a 12% SDS-PAGE (sodium dodecyl sulfate– polyacrylamide gel electrophoresis) with a voltage 10-20 V per cm gel length. The proteins were transferred to PVDF (polyvinylidene difluoride) membranes (0.2 μm pore size, Millipore) by immersion transfer (Bio-Rad). The membrane was then blocked with PBS containing 0.05% Tween 20 and 5% (wt/vol) skim milk, followed by overnight incubation with the anti-Spz^C106^ antibody raised in rat (1:500, a gift from Satoshi Goto, Rikkyo University, Tokyo). Rabbit anti-tubulin (1:1000, Invitrogen) was used as internal control. The primary antibodies were detected with HRP-conjugated anti-rat or anti-rabbit antibodies (Jackson Immunoresearch) and signals were visualized using the Supersignal West Pico Chemiluminescent Substrate (Thermofisher) under ChemiDoc imaging system (Bio-Rad).

### Phenol oxidase (PO) activity assay

Larval hemolymph protein from relevant genotypes were collected as described for western blot. 10 μg of protein mixed with 40 μl of PBS along with the protease inhibitor cocktail in 96 well plate in triplicates and 120 μl of freshly prepared Levodopa (L-DOPA) was added before taking absorbance at 490 nm (see, Ligoxygakis et al., 2002).

### RNA collection and real-time PCR

Hundred developmentally-synchronized larvae of the appropriate genotypes were collected for RNA extraction (Qiagen RNeasy columns). RNA was treated with RNase free DNase, before cDNA synthesis from 2 μg of total RNA. Real time PCR was performed using SYBR green from Applied Biosystems (ABI) on ABI7 900 HT. 100 ng of cDNA sample was mixed with SYBR Green Supermix and appropriate primers to set up a 20 μl reaction mixin triplicates. Transcript levels detected were normalized to actin mRNA values.

Primers used:

*drosomycin:*

Forward primer (5’ gccctcttcgctgtcctga 3’)
Reverse primer (5’ tccctcctccttgcacacac 3’)

### *lgl* and *scrib* tumor transcriptome analysis using Affymmetrix microarray platform

#### Tumor sample collection and RNA isolation

Age-synchronized homozygous larvae of desired genotype (*w*^*118*^ (Control) and *lgl*^*4*^/*lgl*^*4*^ or *scrib*^*vartul*^/*scrib*^*vartul*^ (tumors) were dissected in RNAase free 1X PBS. Wing imaginal discs or their neoplasms were thus obtained were resuspended in 150-300 μL of RLT Buffer (Qiagen) containing 1.5-3 μL of ◻-mercaptoethanol. Total RNA from 30-50 pairs of imaginal discs or tumors were used to make a single sample, which depending on the tumor stage (sizes) typically yields ranging from 10 μg to 15 μg of RNA which were check for quality control and immediately used for labeling and hybridization reactions, using standard protocol recommended by Affymetrix (also, see Khan et al., 2013).

#### Sample Preparation for Affymmetrix Microarray Hybridization

1 μg of RNA was used for two-step cDNA synthesis, involving a first step, T7 Oligo dT Primer-mediated first strand cDNA synthesis (SuperScript II from Invitrogen) followed by second strand synthesis using *E. coli* DNA Polymerase. cDNA was then used to prepare Biotin-labeled RNA by an *in vitro* transcription using IVT labeling kit from Affymetrix Inc, which was fragmented using the Affymetrix supplied Fragmentation buffer (5X = 200 mM Tris-Acetate, pH 8.1, 500 mM KOAc, 150 mM MgOA) and subsequently hybridized to Affymetrix Drosophila Genome 2.0 GeneChip Array.

#### Array Hybridization and Scanning

Hybridization was carried out following manufacturer’s recommendation. 130 μL of hybridization reaction mixture (containing fragmented labeled cRNA, control oligos, hybridization controls, Herring Sperm DNA, Acetylated Bovine Serum Albumin (50 mg/mL) and 2X hybridization buffer) was added to pre-equilibrated array cartridge and the hybridization was allowed to proceed for 16 hours at 45°C. The hybridized arrays were washed and stained using Streptavidin-Phycoerythrin conjugate, and Biotinylated Anti-Streptavidin antibody (Molecular Probes Inc.) to increase the sensitivity. The hybridized arrays were on the Affymetrix microarray scanner.

#### Quality Control, Preprocessing and Data Analysis

Raw data was subjected to quality check using GCOS software, hybridization efficiency was ascertained using spiked B2 Oligo probes, Poly-A RNA controls and *bioB, bioC, bioD* and *cre* labeled controls present in hybridization mix.. The raw .CEL files obtained from GCOS software have been submitted to ArrayExpress (Accession ID: E-MTAB-10059). The .CEL files were analyzed further for differential gene expression using the GenePattern suites (v3.9_20.12.23 build 307) (https://www.genepattern.org/) (Reich et al., 2006). Expression-File-Creator (v13) module was used to create gene expression dataset using RMA (Robust Multi-array Average) algorithm that carries out background correction, quantile normalization, and probe summarization (Irizarry et al., 2003; Li, 2001). Comparative-Marker-Selection (v10) module was used to compute fold changes and two-sided *t*-test was used as a statistical measure of significance. List of Toll-related genes was manually computed. Dendogram representing agglomerative hierarchical clustering for genes (rows) and phenotypes (columns) was computed for select genes from the larger Toll pathway gene set using Hierarchical-clustering (V6) module (Eisen et al., 1998). Pearson correlation was used for row distance measure while pairwise average linkage was used to compute cluster distances. Dendogram represented in the computed Gene tree file was used as an input into Hierarchical Cluster Image module (v4) to generate heatmaps, with row normalized color scheme. Labels of heatmaps were further edited in Adobe Photoshop (CS6) for clarity of fonts.

#### Gene Set Enrichment Analysis (GSEA)

Gene-set enrichment analysis provides a quantitative measure of the enrichment of predefined gene-sets between two phenotypes being compared (Mootha et al., 2003; Subramanian et al., 2005). A normalized enrichment score (NES) is calculated using running sum statistics, which is a positive value if positively correlated with the phenotype under study (in this case tumor (*lgl* or *scrib*) versus control (wild type). Raw .CEL files for individual Affymmetrix run were pre-processed for background correction, quantile normalization, using GenePattern (https://www.genepattern.org/), and the .gct file served as an input for the GSEA. Pre-defined gene set for Toll signaling pathway was manually computed. GSEA analyses were carried out using the following parameters: genes were ranked based on comparison of phenotypes: *lgl* or *scrib* (test) vs. control (wild type) using signal-to-noise metric. The enrichment score was calculated using weighted running sum statistics. Gene-based permutation (n = 1,000) was used to compute the nominal P value and FDR was computed to correct for multiple-hypothesis testing.

## Graphical Summary

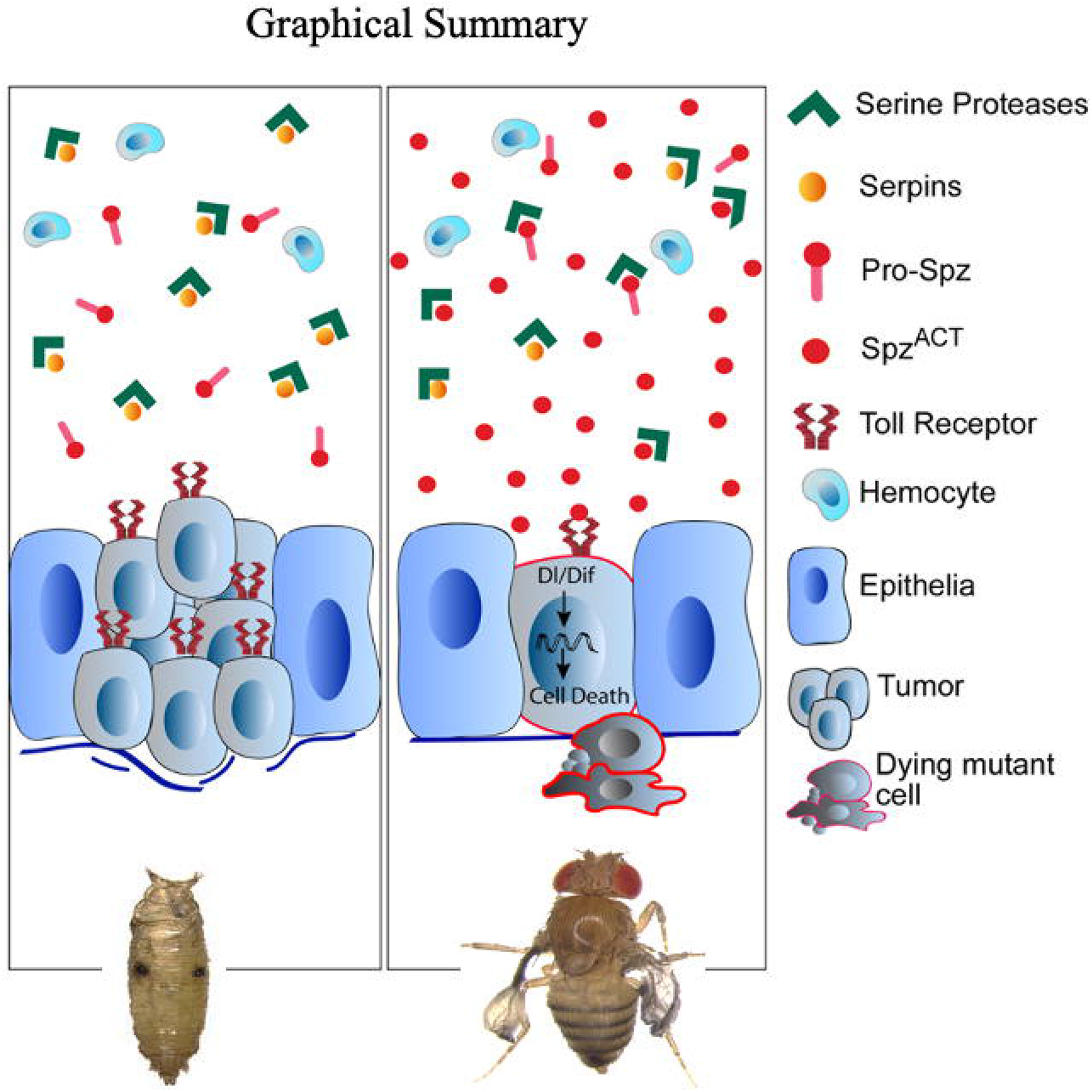

### Highlights

- *Drosophila* neoplasm-derived Spz^Act^ ligand triggers both localized and systemic host Toll signaling
- Tumor-trigger Toll signaling, however, does not suffice for tumor elimination
- A *spn27A*^*1*^/+ host genetic background displays robust and constitutive host Toll signaling
- Prophylactic host Toll signaling brings about NF-KB-mediated tumor remission

### In Brief

In *Drosophila*, a host genetic background displaying heterozygosity of *serpin 27A*^*1*^ mutation displays constitutive Toll signaling and confers cancer resistance via NF-KB-mediated tumor cell death.

